# Vimentin is Required for Tumor Progression and Metastasis in a Mouse Model of Non-Small Cell Lung Cancer

**DOI:** 10.1101/2020.06.04.130963

**Authors:** Alexandra L. Berr, Kristin Wiese, Gimena dos Santos, Jennifer M. Davis, Clarissa M. Koch, Kishore R. Anekalla, Martha Kidd, Yuan Cheng, Yuan-Shih Hu, Karen M. Ridge

## Abstract

Vimentin, a type III intermediate filament, is highly expressed in aggressive epithelial cancers and is associated with increased rates of metastasis. We show that vimentin is causally required for lung cancer metastasis using a genetic mouse model of lung adenocarcinoma (*LSL*-*Kras*^G12D^;*Tp53*^fl/fl^, termed *KPV*^+/+^) crossed with vimentin-null mice (thereby creating *KPV*^−/−^ mice). Both *KPV*^+/+^ and *KPV*^−/−^ mice developed lung tumors, yet *KPV*^−/−^ mice had delayed tumorigenesis and prolonged survival. *KPV*^+/+^ cells implanted in the flank metastasized to the lung while *KPV*^−/−^ cells did not, providing additional evidence that vimentin is required for metastasis. Differential expression analysis of RNA-seq data demonstrated that *KPV*^−/−^ cells had suppressed expression of genes that drive epithelial-to-mesenchymal transition, migration, and invasion, processes that are critical to the metastatic cascade. Integrative metabolomic and transcriptomic analysis revealed altered glutaminolysis, with *KPV*^−/−^ cells accumulating glutathione, leading to impaired cell motility in response to oxidative stress. Together, these results show that loss of vimentin impairs epithelial-to-mesenchymal transition and regulation of the oxidative stress response, resulting in decreased metastasis in murine lung adenocarcinoma.

## Introduction

Non-small-cell lung cancers (NSCLCs) represent 80% of all lung cancers and are often diagnosed at more advanced stages of the disease resulting in high rates of mortality (1). Adenocarcinoma is the most common subtype of NSCLC and is characterized by activating mutations in the *Kras* proto-oncogene in up to 30% of diagnoses and by inactivating mutations in the tumor suppressor gene *Tp53* in up to 60% of diagnoses (2–5). Despite the prevalence of lung adenocarcinoma, the metastatic mechanisms that drive lung cancer progression are incompletely understood.

The type III intermediate filament vimentin is associated with increased metastatic spread and lower rates of survival in patients with NSCLC (6–8). Vimentin is a canonical marker of epithelial-to-mesenchymal transition (EMT), an initiating event of the metastatic cascade (9). EMT is the process by which epithelial cells remodel cell-cell and cell-extracellular-matrix (ECM) contacts, lose their apical-basal polarity, and adopt the spindle-shaped morphology associated with a mesenchymal cell phenotype (10). During EMT, cells undergo a downregulation of epithelial cell associated genes, including E-cadherin and cytokeratins, and an upregulation of mesenchymal cell associated genes, including N-cadherin and vimentin. In addition to acting as a marker of EMT, vimentin is functionally involved in EMT. Structurally, vimentin intermediate filaments control cell shape, and thus facilitate the transition toward a mesenchymal phenotype (11). Twist1, a transcription factor critical to EMT, upregulates vimentin expression (12). Vimentin also serves as a scaffold for Slug, another transcription factor that regulates EMT (13). Ultimately, EMT leads to an increase in epithelial-derived cell motility, which marks the first step in cancer metastasis. Our laboratory has previously shown that vimentin is required for the EMT-like process responsible for epithelial wound repair (14). In addition to its role in EMT, vimentin cooperates with actin and microtubules to mediate invasion across the basement membrane and migration through the collagen-rich interstitial space. Cancer cells form invadopodia, proteolytically active plasma membrane projections that break down the basement membrane (15). Through an indirect interaction with actin, vimentin intermediate filaments participate in the elongation of invadopodia, which allows cells to traverse the basement membrane and escape from the primary tumor site toward the nearest capillary. As a cell begins to migrate, vimentin is crucial for the development of cellular polarity, which is necessary for the efficient migration of tumor cells (16). This cancer cell migration is mediated through activation of the PI3K/Akt pathway. Within this signaling cascade, Akt1 phosphorylates vimentin, which leads to downstream increases in cell motility by protecting vimentin filaments from proteolysis (17). When the PI3K/Akt pathway is blocked, vimentin expression is attenuated; this decrease in vimentin is associated with a lower rate of pulmonary metastases in a murine breast cancer model (18). Together, these mechanisms are responsible for a decrease in migration, invasion, and metastasis conferred by a loss of vimentin in lung, breast, head and neck, and bone cancer cells (11, 17–24).

Although numerous studies have provided correlative data and *in vitro* data to establish the link between vimentin and aspects of the metastatic cascade, none have provided evidence to suggest that vimentin plays a causal role in NSCLC metastasis. Therefore, we set out to characterize the role of vimentin in NSCLC metastasis using a genetically engineered mouse model (GEMM). In the present study, we used the well-established *LSL*-*Kras*^G12D^;*Tp53*^fl/fl^ (*KPV*^+/+^) mouse model, which reliably recapitulates human NSCLC in pathology, disease progression, clinical outcome, and response to therapies (25). To identify the role of vimentin in lung adenocarcinoma, we crossed this GEMM with the global vimentin knockout (*Vim^−/−^*), thereby creating *KPV*^−/−^ mice (26).

In this study, we show that *KPV*^−/−^ mice have reduced lung tumor burden and increased rates of survival compared to *KPV*^+/+^ mice. Of note, *KPV*^−/−^ cells have significantly impaired metastatic potential. Mechanistically, we find that *KPV*^−/−^ cells display impaired EMT; RNA sequencing (RNA-seq) and cell motility assays reveal that *KPV*^−/−^ cells fail to adopt a mesenchymal phenotype while *KPV*^+/+^ cells do. Our data suggest that the targeted loss of vimentin may serve as a therapeutic strategy by which to disrupt the development of lung adenocarcinoma and suppress metastasis.

## Results

### Generation of an LSL-Kras^G12D/+^; Tp53^fl/fl^;Vim^−/−^ (KPV^−/−^) genetically engineered mouse model

Tumorigenesis and metastasis are well characterized in the *LSL-Kras*^G12D/+^;*Tp53^fl/fl^* (*KPV*^+/+^) GEMM (25). When adenoviral Cre recombinase (Ad-Cre) is delivered intratracheally to *KPV*^+/+^ mice, tumors develop as early as 2 weeks post-infection (wpi) (25). This model recapitulates the highly invasive nature of lung adenocarcinoma with ~50% of mice developing metastatic lesions in the mediastinal lymph nodes and the pleural spaces of the thoracic cavity (27). We crossed a global vimentin knockout mouse to the *KPV*^+/+^ mouse to create the *KPV*^−/−^ mouse (26) (***SI Appendix***, **S1A**). This novel *KPV*^−/−^ mouse lacks vimentin expression throughout the lungs at baseline (***SI Appendix***, **S1B**). Following Ad-Cre administration, disruption of the *Kras* allele and accumulation of mutant KRAS protein was validated by reverse transcription polymerase chain reaction (RT-PCR) and Western blot, respectively (***SI Appendix***, **S1C-D**). *Rosa26-LSL-LacZ* reporter mice were used to validate the intratracheal delivery of Ad-Cre (28). Mice infected with Ad-Cre demonstrated homogenous, positive lacZ expression, while mice treated with adenoviral null construct (Ad-null) did not express lacZ (***SI Appendix***, **S1E**). These results demonstrate the utility of a novel *KPV*^−/−^ GEMM to define a causal role of vimentin in lung adenocarcinoma metastasis.

### Vimentin deficiency increases survival and reduces tumor burden

Weight loss, also termed “cancer cachexia,” is a common manifestation of morbidity in human cancer patients and is associated with a poor prognosis in patients with advanced disease. *KPV*^+/+^ and *KPV*^−/−^ mice were administered Ad-Cre and their weight was recorded weekly. *KPV*^+/+^ mice showed a rapid and profound decline in total body weight starting at 4 wpi, while *KPV*^−/−^ mice did not exhibit weight loss until 9 wpi, suggesting less advanced disease in the vimentin-deficient mice (**Figure 1A**). *KPV*^−/−^ mice lived significantly longer than *KPV*^+/+^ mice, with a median survival of 15.5 wpi compared to 10 wpi in the *KPV*^+/+^ mice (**Figure 1B**). Lung tumor development was confirmed in *KPV*^+/+^ and *KPV*^−/−^ mice using magnetic resonance imaging (MRI). At 6 wpi, *KPV*^−/−^ mice had an average lung tumor burden of 7.5%, which was significantly lower than that of 37% seen in *KPV*^+/+^ mice (**Figure 1C**, ***SI Appendix***, **S2A**). Lungs were harvested, fixed, and stained with hematoxylin and eosin (H&E). *KPV*^+/+^ mice displayed a greater increase in hyperplastic lesions at 8 wpi (36 ± 5 hyperplasias per *KPV*^+/+^ mouse vs. 14 ± 2 hyperplasias per *KPV*^−/−^ mouse). At 12 wpi, *KPV*^+/+^ mice displayed increased numbers of both adenomas and adenocarcinomas (6.3 ± 1.8 adenomas and 1.2 ± 0.2 adenocarcinomas per *KPV*^+/+^ mouse) compared to *KPV*^−/−^ mice (1.6 ± 1.0 adenomas and 0.04 ± 0.02 adenocarcinomas per *KPV*^−/−^ mouse) (***SI Appendix***, **S2B**). Together, these data suggest that tumor progression is suppressed by the loss of vimentin in *KPV*^−/−^ mice. Immunohistochemistry (IHC) staining for vimentin, TTF-1, and Ki67 was performed on serial sections of lung tissue from *KPV*^+/+^ and *KPV*^−/−^ mice at 6 wpi. Vimentin was expressed in *KPV*^+/+^ tumors and normal adjacent lung tissue, but was not expressed in *KPV*^−/−^ lung tissue (**Figure 1D**). TTF-1 is a biomarker associated with lung adenocarcinoma (29). *KPV*^−/−^ mice had fewer TTF-1-positive cells, and clusters of TTF-1-positive cells were smaller than those observed in *KPV*^+/+^ lungs. *KPV*^+/+^ and *KPV*^−/−^ mice displayed positive Ki67 staining, which was mainly associated with tumor cells, with no apparent difference in quantity or localization of the dividing cells as determined by Ki67 staining (**Figure 1E**). Mutant KRAS activates the RAF–MEK–ERK pathway, which is involved in cancer cell proliferation and survival (30). IHC revealed similar levels of pERK1/2 staining between *KPV*^+/+^ and *KPV*^−/−^ mice, suggesting that mutant KRAS expression was activated in *KPV*^+/+^ and *KPV*^−/−^ mice (***SI Appendix***, **S2C**). Because infiltrating immune cells can serve as tumor suppressors or promotors, we evaluated the presence of immune cells by IHC staining with an antibody against CD45 (***SI Appendix***, **S2C**). Strikingly, lung tissue from *KPV*^−/−^ mice had fewer CD45-positive stained cells than did *KPV*^+/+^ mice. Finally, by 12 wpi, *KPV*^+/+^ mice accumulated mutant *Kras* transcripts in the liver while *KPV*^−/−^ mice did not, suggesting that vimentin-expressing cells form metastatic lesions (**Supplemental Figure 1C**). Together, these data suggest that loss of vimentin suppresses tumor development in this mouse model of lung adenocarcinoma, which leads to prolonged survival.

**Figure 1.**
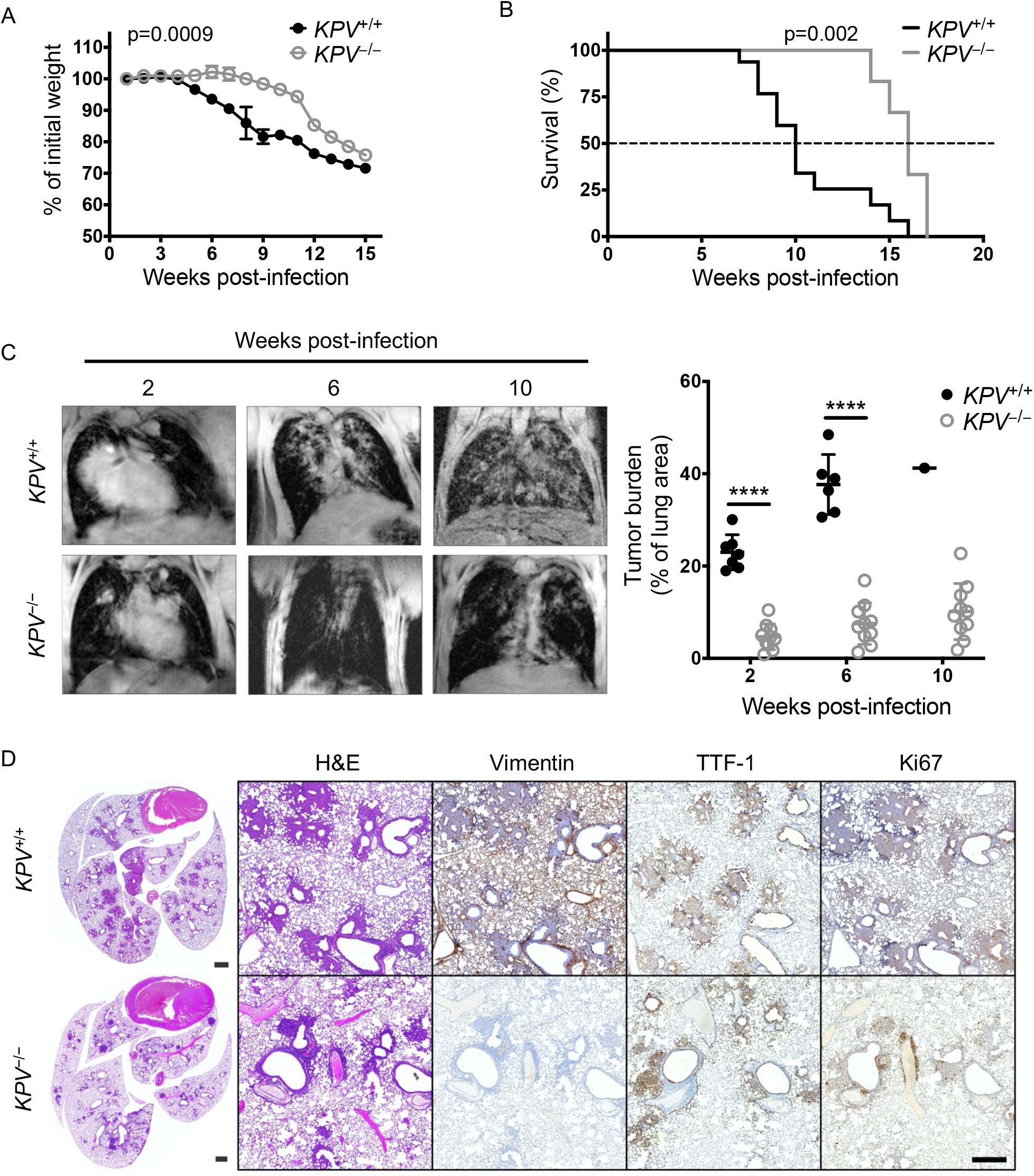
Vimentin-null mice have reduced tumor burden and improved survival in a preclinical *LSL-Kras*^*G12D*^*Tp53*^*fl/fl*^-driven mouse model of lung cancer. *LSL*-*Kras*^*G12D*^*Tp53*^*fl/fl*^ (*KPV*^+/+^) mice were crossed with *Vim*^−/−^ mice to produce *KPV*^−/−^ mice, then *KPV*^+/+^ and *KPV*^−/−^ mice were intubated with 109 PFUs of adenoviral Cre. Weight loss (n=6 mice for *KPV*^+/+^ group; n=7 mice for KPV^−/−^ group; mixed model ANOVA, for *KPV*^+/+^ versus *KPV*^−/−^, p=0.0009) and (**B**) survival (n=15 mice for *KPV*^+/+^ group; n=10 mice for *KPV*^−/−^ group; Mantel-Cox log-rank test, p=0.002) were monitored. (**C**) Representative MRI scans (*left*) showing mouse lung tumors at 2, 6, and 10 weeks post-infection with 109 PFUs of adenoviral Cre. Tumor burden was quantified using Jim software (*right*). Each point represents one mouse (****p<0.0001 by unpaired, two-tailed t-test). (**D**) Lungs were isolated from *KPV*^+/+^ mice (6 weeks post-infection shown) and *KPV*^−/−^ mice (7 weeks post-infection shown) infected with 109 PFUs of adenoviral Cre. Shown from left to right are representative fixed whole lung sections with H&E staining and close-up views of fixed lung sections with H&E staining and vimentin, TTF-1, and Ki67 immunohistochemical staining. Positively immunostained cells appear brown, and nuclei are dyed blue. Scale bars: 1 mm (whole lungs, *left*), 200 μm (*right*). This figure represents combined data from three independent experiments.

### Transcriptional profiling reveals a less differentiated cancer phenotype in vimentin-null cancer cells

To better understand how vimentin is involved in the molecular pathways of lung adenocarcinoma, we used RNA-seq to identify genes that have altered expression in the absence of vimentin. To this end, epithelial-derived cancer cells (CD45-negative, EpCAM-positive) were isolated from *KPV*^+/+^ and *KPV*^−/−^ lungs at 6 wpi with Ad-Cre (hereafter referred to as *KPV*^+/+^ and *KPV*^−/−^ cells). The absence of vimentin in *KPV*^−/−^ cells was confirmed via Western blot and immunofluorescence staining (***SI Appendix***, **S1F and G**). We isolated messenger RNA (mRNA) from *KPV*^+/+^ and *KPV*^−/−^ cells and performed RNA-seq. Samples clustered together via principal component analysis (PCA) and Pearson’s correlation with no outlying data points (***SI Appendix***, **S3A and B**). A majority of the sample variance (97.6%) could be attributed to the loss of vimentin in *KPV*^−/−^ cells (***SI Appendix***, **S3A**). There were 904 differentially expressed genes (DEGs) between the *KPV*^+/+^ and *KPV*^−/−^ cells (***SI Appendix***, **S3C**). Of these, 316 genes were upregulated (Cluster 1) and 588 genes were downregulated (Cluster 2) in *KPV*^+/+^ cells compared to *KPV*^−/−^ cells (**Figure 2A**). To characterize these genes, we performed Gene Ontology (GO) enrichment. Of note, epithelial cell differentiation and cell adhesion genes were upregulated in *KPV*^−/−^ cells while cell migration and mesenchymal cell proliferation genes were downregulated in *KPV*^−/−^ cells. When we explored the DEGs that contribute to these pathways, we found that several EMT-associated genes are upregulated in *KPV*^+/+^ cells. These include *Twist1* and *Cdh2*, the gene that codes for N-cadherin (**Figure 2B**). This finding was confirmed by Western blot, which showed an increase in N-cadherin in *KPV*^+/+^ cells compared to *KPV*^−/−^ cells (***SI Appendix***, **1F**). In contrast, genes associated with epithelial cell phenotype, including claudins *Cldn2, Cldn8*, and *Cldn18*, as well as the cytokeratins *Krt4, Krt20*, and *Krt23*, were upregulated in *KPV*^−/−^ cells. These data suggest that *KPV*^−/−^ cells retain the phenotype associated with their alveolar epithelial cell origin and fail to upregulate key mesenchymal genes that confer metastatic potential. Therefore, *KPV*^−/−^ cells lack key mesenchymal features that *KPV*^+/+^ cells adopt. Invasion is potentiated by matrix metalloproteases (MMPs), which break down the basement membrane, allowing cells to move toward adjacent capillaries. *Mmp11, Mmp15*, and *Mmp24* are upregulated in *KPV*^+/+^ cells (**Figure 2B**). To cross the basement membrane, cells must form invadopodia, a process that relies on vimentin (15). Accordingly, invadopodia-associated genes (*Arpc2, Arpc5, Arpc5l*, and *Actr2*) are downregulated in *KPV*^−/−^ cells (**Figure 2B**). Cells must then migrate across a collagen-rich interstitial space to reach the bloodstream. This process is coordinated by chemokines (*Cxcl12*), integrins (*Itga5* and *Itgb5*), and alterations in the ECM (*Lama3, Lamb3, Lamc2*, and *Fn1*); these genes are significantly downregulated in *KPV*^−/−^ cells compared to *KPV*^+/+^ cells (**Figure 2B**). Together, these results suggest that vimentin is involved in the early cellular events that lead to metastasis.

**Figure 2.**
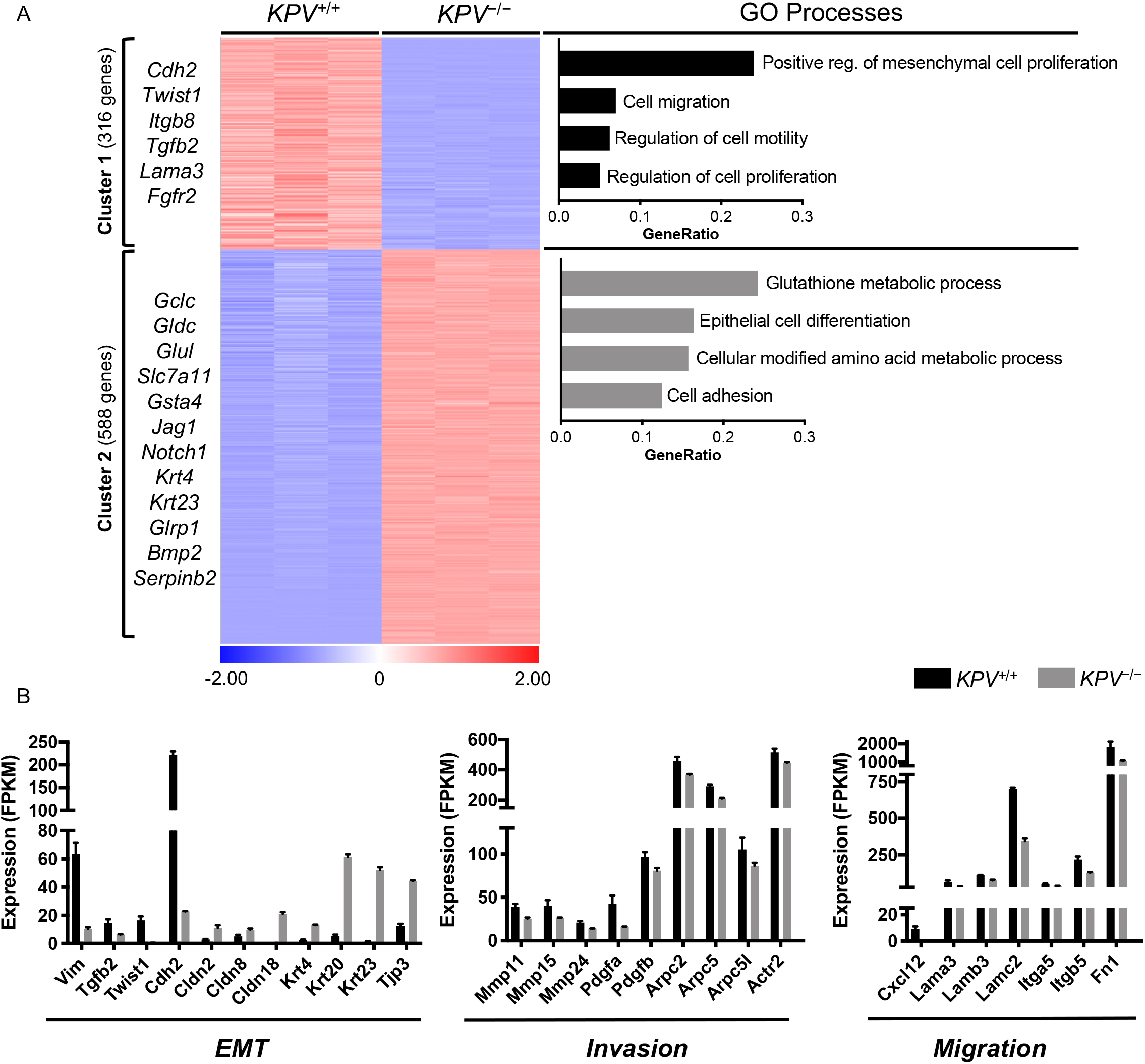
*KPV*−/−cells have decreased expression of genes involved in EMT. Messenger RNA collected from *KPV*^+/+^ and *KPV*^−/−^ cells was subjected to RNA-sequencing. (**A**) Differentially expressed genes (DEGs) between *KPV*^+/+^ and *KPV*^−/−^cells were clustered using K-means clustering. Genes enriched in Cluster 1 (316 genes) and Cluster 2 (588 genes) were subject to GO enrichment analysis. GO Processes with FDR<0.05 are shown. GeneRatio is the number of genes present in the cluster that are associated with the GO process divided by the total number of genes in that GO process. (**B**) Expression values (FPKM) of select genes are shown. N=3 for each group. Data in panel **B** are presented as the mean ± standard deviation. All gene comparisons shown (*KPV*^+/+^ vs *KPV*^−/−^) have FDR<0.05 after adjusting for multiple comparisons.

### An intact vimentin network is required for cancer cell migration and invasion

Based on the observation that cell motility pathways are downregulated in *KPV*^−/−^ cells compared to *KPV*^+/+^ cells, we set out to test whether vimentin is required for the cell-intrinsic motility properties that are necessary for lung cancer cell metastasis. *KPV*^+/+^ and *KPV*^−/−^ cells were subjected to several motility assays that mimic the transit patterns a lung cancer cell undergoes during metastasis. To evaluate migration, a scratch wound healing assay was performed (**Figure 3A**). Within 6 hours, *KPV*^+/+^ cells had closed the majority of the wound area (72.71 ± 3.267%). In contrast, *KPV*^−/−^ cells closed only 17.71 ± 3.267% of the denuded area. To test the invasive potential of *KPV*^+/+^ and *KPV*^−/−^ cells, a Matrigel-coated transwell assay was used to mimic invasion across the alveolar basement membrane. *KPV*^+/+^ cells had a 16-fold increased rate of invasion as compared to *KPV*^−/−^ cells (invasive index, 230 ± 41.76 vs. 14.58 ± 2.68, respectively) (**Figure 3B**). Three-dimensional invasion was modeled by generating *KPV*^+/+^ and *KPV*^−/−^ spheroids and tracking the invasion of cells through collagen gels. *KPV*^+/+^ spheroids grew 4.65 times larger than *KPV*^−/−^ spheroids, suggesting that, in a three-dimensional model, *KPV*^−/−^ cells have impaired migration and invasion (**Figure 3C**). Given these results, we conclude that vimentin is required for migration and invasion in this model of lung adenocarcinoma.

**Figure 3.**
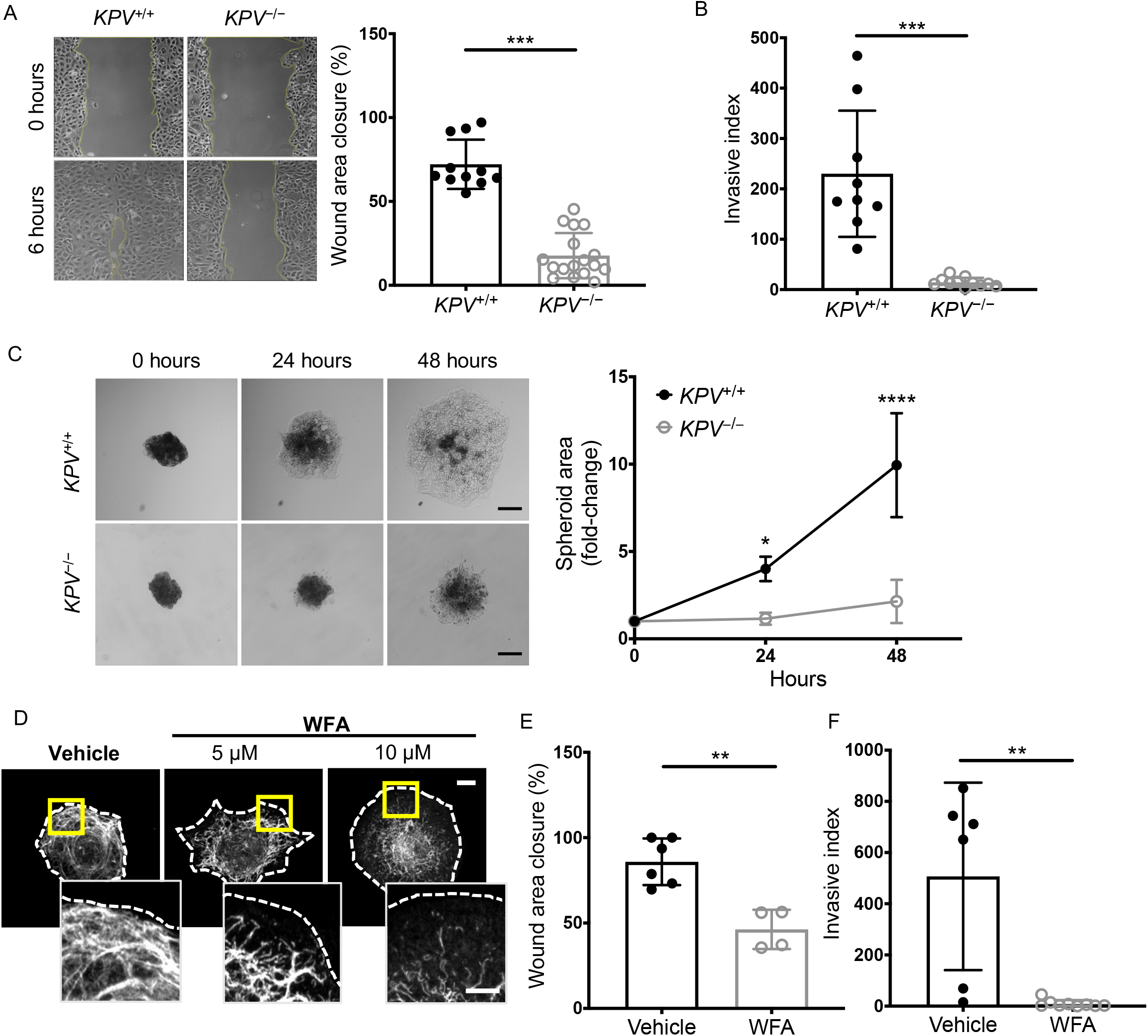
Vimentin is required for *in vitro* cancer cell migration and invasion. **(A)** A scratch wound assay was used to evaluate cell migration. Representative images are shown at 0 and 6 hours following scratch formation. Wound area closure was compared to the starting value and quantified for *KPV*^+/+^ (n=11) and *KPV*^−/−^ (n=17) cells; each point represents a separate scratch wound. **(B)** Cell invasion through a Matrigel-coated transwell was measured over 48 hours. Invasive index is the mean number of cells invaded per 20× magnification imaging field. *KPV*^+/+^ (n=9) and *KPV*^−/−^ (n=11) cell invasion data are plotted so that each point represents data from a single transwell assay. **(C)** *KPV*^+/+^ and *KPV*^−/−^ spheroids were suspended in type I collagen and spheroid growth was tracked over 48 hours. Spheroid area was quantified relative to the initial area of each spheroid (n=4 independent experiments). Scale bar: 200 μM. **(D)** *KPV*^+/+^ cells were treated with withaferin A (WFA; 5 or 10 μM) or DMSO vehicle control for 6 hours. Cells were stained for vimentin (white). Cell outline (dotted line) was drawn on an image of the same cell stained for keratin. Scale: 10 μm; inset: 5 μm. **(E)** *KPV*^+/+^ cells were treated with vehicle (n=6) or 5 μM WFA (n=4) and were subjected to a scratch wound assay. Wound area was quantified at 6 hours. (F) KPV^+/+^ cells were plated atop a Matrigel-coated transwell and were treated with vehicle control (n=6) or 5 μM WFA (n=9); invasion was quantified at 48 hours via invasive index as described above. Data are presented as the mean ± standard deviation. The p-values were calculated using an unpaired, two-tailed t-test, except for panel C, in which data were compared using a repeated-measure two-way ANOVA with multiple comparisons. (*p<0.05; **p<0.01; ***p<0.001; ****p<0.0001).

Withaferin A (WFA) is a steroidal lactone that has been validated as an anticancer agent in a range of murine tumor models, including breast, prostate, and ovarian cancer (31–33). WFA causes vimentin aggregation in cells by promoting phosphorylation at serine 38 (Ser38) and serine 56 (Ser56) (34). We set out to determine whether WFA-mediated disruption of the vimentin intermediate filament network affects lung adenocarcinoma cell metastasis. To test the hypothesis that the vimentin intermediate filament network is sufficient for migration and invasion *in vitro*, we treated *KPV*^+/+^ cells and human lung adenocarcinoma (A549) cells with WFA. The vimentin intermediate filament network, which extends from the nucleus to the plasma membrane, was retracted from the plasma membrane and collapsed around the nucleus following treatment with WFA, with no change in vimentin protein expression (**Figure 3D**, **Supplemental S4A-B**) (35). WFA treatment decreased *KPV*^+/+^ migration in a scratch wound assay by ~46% (**Figure 3F**). Similarly, cell invasion was completely suppressed following treatment with WFA (**Figure 3G**). We observed a similar trend in human-derived A549 cells, which exhibited a dose-dependent decrease in cell migration following treatment with WFA (***SI Appendix***, **S3C**). These data suggest that disruption of the vimentin intermediate filament network impairs lung adenocarcinoma cell motility.

### Withaferin A treatment attenuates cancer progression

We set out to determine whether WFA-mediated disruption of the vimentin intermediate filament network affects lung adenocarcinoma progression. Mice were administered Ad-Cre to initiate tumor development; at 2 wpi, *KPV*^+/+^ mice were administered WFA (4 mg/kg, Q.O.D, p.o.) (**Figure 4A**). At 6 wpi, *KPV*^+/+^ mice that were given WFA had developed smaller tumors (tumor burden, 15.65 ± 2.518%) than had vehicle-treated mice (tumor burden, 25.1 ± 3.842%) (**Figure 4B**). Lungs were harvested, fixed, and stained with H&E and immunostained for vimentin, TTF-1, and Ki67. Vehicle-treated *KPV*^+/+^ mice had enhanced TTF-1 and Ki67 staining associated with lung tumors. In contrast, WFA-treated mice had reduced tumor burden with diminished TTF-1 and Ki67 staining (**Figure 4C**). Collectively, these data suggest that vimentin can be pharmacologically targeted to disrupt the ability of lung cancer cells to invade and migrate away from the primary tumor.

**Figure 4.**
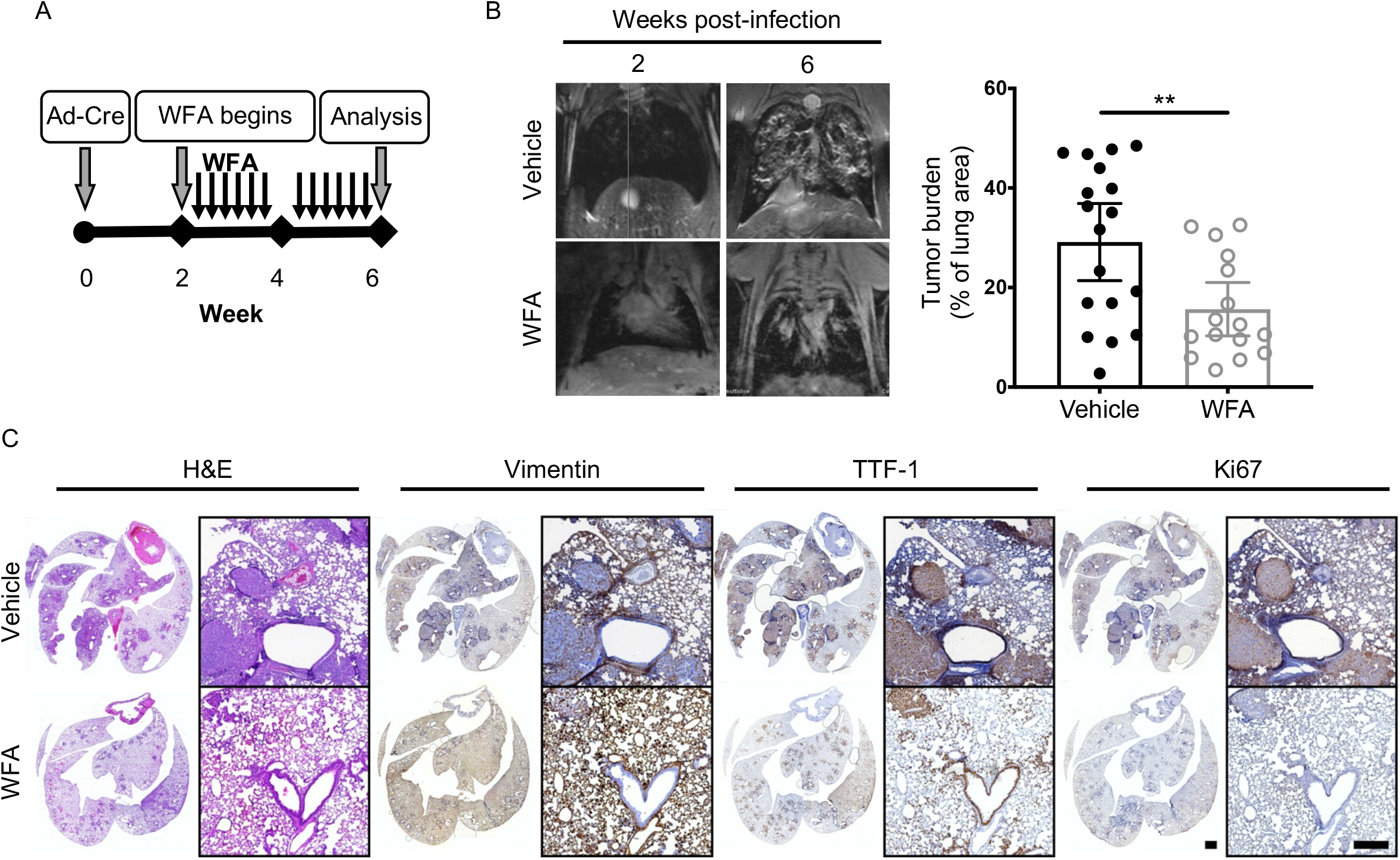
WFA treatment attenuates lung cancer progression. (**A**) Schematic of experimental design. *KPV*^+/+^ mice were treated with withaferin A (WFA; 4 mg/kg; Q.O.D., p.o.) or vehicle control (DMSO) at 2 weeks post-infection with 107 PFUs of adenoviral Cre. (**B**) Representative MRI scans show WFA-treated *KPV*^+/+^ lung tumors at 6 weeks post-infection with 107 PFUs of adenoviral Cre (*left*). Dot plot illustrates the tumor volume between WFA-treated or vehicle-treated control *KPV*^+/+^ mice (*right*). Each point represents, for one mouse, the percentage of lung area on MRI occupied by tumor, as measured using Jim software. Data are presented as the mean ± standard deviation (**p<0.01 by unpaired, two-tailed t-test). (**C**) Lungs isolated from vehicle- or WFA-treated KPV^+/+^ mice at 6 weeks after adenoviral Cre infection were fixed, sectioned, and subjected to H&E staining and vimentin, TTF-1, and Ki67 immunohistochemical staining. Positively immunostained cells appear brown, and nuclei are dyed blue. Scale bars: 2 mm (whole lungs, *left*), 200 μM (*right*).

### Vimentin-null cancer cells accumulate glutathione

Glutathione is an antioxidant that neutralizes reactive oxygen species (ROS) and thus regulates cellular response to oxidative stress. We identified genes involved in glutathione and serine metabolic processes that were significantly upregulated in *KPV*^−/−^ cells compared to *KPV*^+/+^ cells (**Figure 2A**). Furthermore, genes associated with glycolysis are positively enriched in *KPV*^−/−^ cells while genes associated with oxidative phosphorylation are negatively enriched in *KPV*^−/−^ cells (***SI Appendix***, **S3D**). To better understand how the loss of vimentin leads to changes in the metabolic landscape, we performed metabolomics analysis on *KPV*^+/+^ and *KPV*^−/−^ cells. In *KPV*^−/−^ cells, glutathione and several metabolites involved in in its production were upregulated compared to *KPV*^+/+^ cells; these other metabolites include serine, glycine, glutamate, glutamine, and cystathionine (**Figure 5A**). By interrogating our RNA-seq data, we identified significantly upregulated genes that correspond to an increased production of glutathione in *KPV*^−/−^ cells. These include: *Slc1a5*, a glutamine transporter; *Shmt1*, an enzyme required for metabolism of serine to glycine; and *Gclm* and *Gclc*, enzymes involved in the conversion of glutamate to glutathione (**Figure 2A**, **Figure 5C)**. The RNA-seq and metabolomics findings are summarized in **Figure 5D** and show that, compared to *KPV*^+/+^ cells, *KPV*^−/−^ cells have elevated levels of metabolites and genes involved in the production of glutathione.

**Figure 5.**
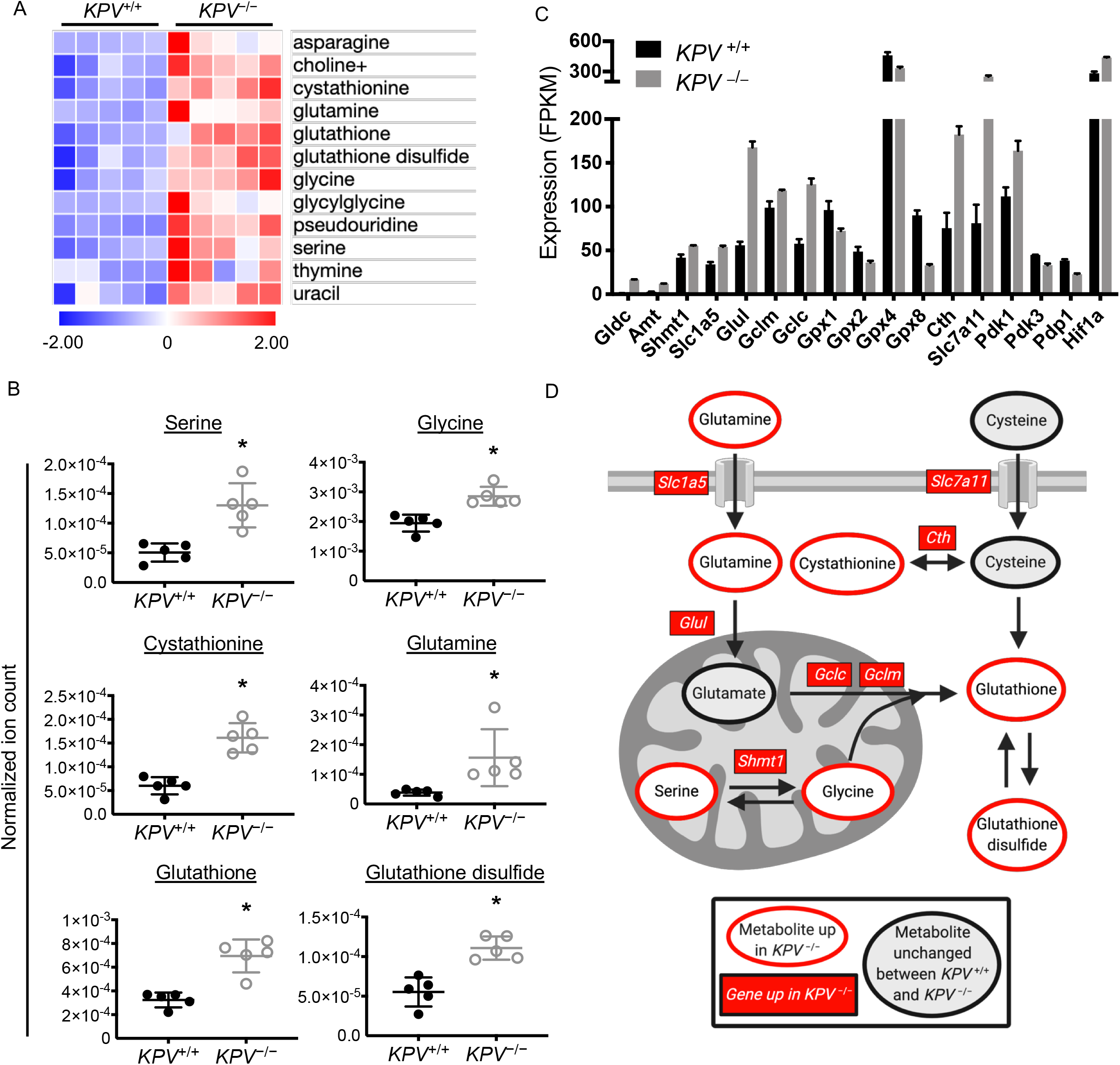
*KPV*^−/−^ cells accumulate glutathione. (**A**) Differentially produced metabolites are plotted with each row representing z-scores for each metabolite. (**B**) Ion counts were normalized to the total ion count for each sample. Each group represents n=5. For panels **A-B**, metabolite data was log-transformed and then subjected to an unpaired two-tailed t-test; p-values were corrected for multiple comparisons (**\*** adjusted p-value<0.05). (**C**) Select gene expression values from RNA-sequencing. All genes shown had FDR<0.05. (**D**) Schematic showing key metabolites (*ovals*) and genes (*rectangles*) involved in cell production of glutathione. Metabolites and genes downregulated in *KPV*^−/−^ cells compared to *KPV*^+/+^ cells are shown in red. All data are presented as the mean ± standard deviation.

### Hypoxia-mediated cell migration is dependent on vimentin

Based on the finding that *KPV*^−/−^ cells produce higher levels of glutathione than do *KPV*^+/+^ cells, we hypothesized that vimentin is involved in the lung adenocarcinoma response to oxidative stress. Vimentin is involved in a ROS negative feedback loop: high levels of ROS increase vimentin expression, and vimentin filaments protect cells from oxidative damage and lead to decreased production of ROS (36–38). Compared to *KPV*^+/+^ cells, *KPV*^−/−^ cells produce high levels of hypoxia-inducible factor 1-alpha (*Hif1a*), the master regulator of the cellular response to hypoxia (**Figure 5C**). Hypoxia occurs when tumors outgrow their blood supply and the overall amount of oxygen available for cancer cell respiration decreases (39). Hypoxic environments cause mitochondria to produce ROS, which can promote EMT and metastasis (40–42). Exposure to hypoxia *in vitro* does not change overall vimentin protein content of A549 cells (**Figure 6A**). However, hypoxia impacts the organization of the vimentin intermediate filament network in a similar manner as WFA treatment (**Figure 6B**). Under normoxic conditions, vimentin filaments extend from the nucleus to the periphery of the cell. In contrast, hypoxia causes retraction of vimentin intermediate filaments from the plasma membrane and formation of disassembled “squiggles” at the cell edge. To test the functional outcomes of this shift in architecture, we subjected wild-type (*Vim*^+/+^) and Vimentin knockdown (*Vim*^KD^) cells to motility assays (**Figure 6C**). Under hypoxia, *Vim*^+/+^ cells have ~1.43 times greater wound closure than under normoxic conditions. *Vim*^KD^ have impaired wound closure under hypoxia compared to normoxia, with a relative migration rate of 0.73 times their rate under normoxic conditions (**Figure 6D**). Similarly, the invasive index of Vim^+/+^ cells is 3.33 times higher under hypoxic conditions compared to normoxic conditions, an increase that was not observed with *Vim*^KD^ cells (**Figure 6E**). For both migration and invasion assays, hypoxia led to significantly higher rates of motility in *KPV*^+/+^ cells compared to *Vim*^KD^ cells. Hypoxia activates the PI3K/Akt pathway (43). Accordingly, we observed an accumulation of phosphorylated Akt (pAkt) over 24 hours of exposure to hypoxia (**Figure 6F**). However, in the absence of vimentin, pAkt levels decreased (**Figure 6G**). To identify whether vimentin and pAkt physically interact, we performed immunoprecipitation on vimentin collected from cells cultured under either normoxia or hypoxia. We found that, under hypoxia, pAkt binds vimentin (**Figure 6H**). These findings are supported by previous reports that Akt1 activation mediates cell invasion in soft-tissue sarcoma through its interaction with vimentin (17). Therefore, we concluded that vimentin is required for hypoxia-mediated cell invasion and migration.

**Figure 6.**
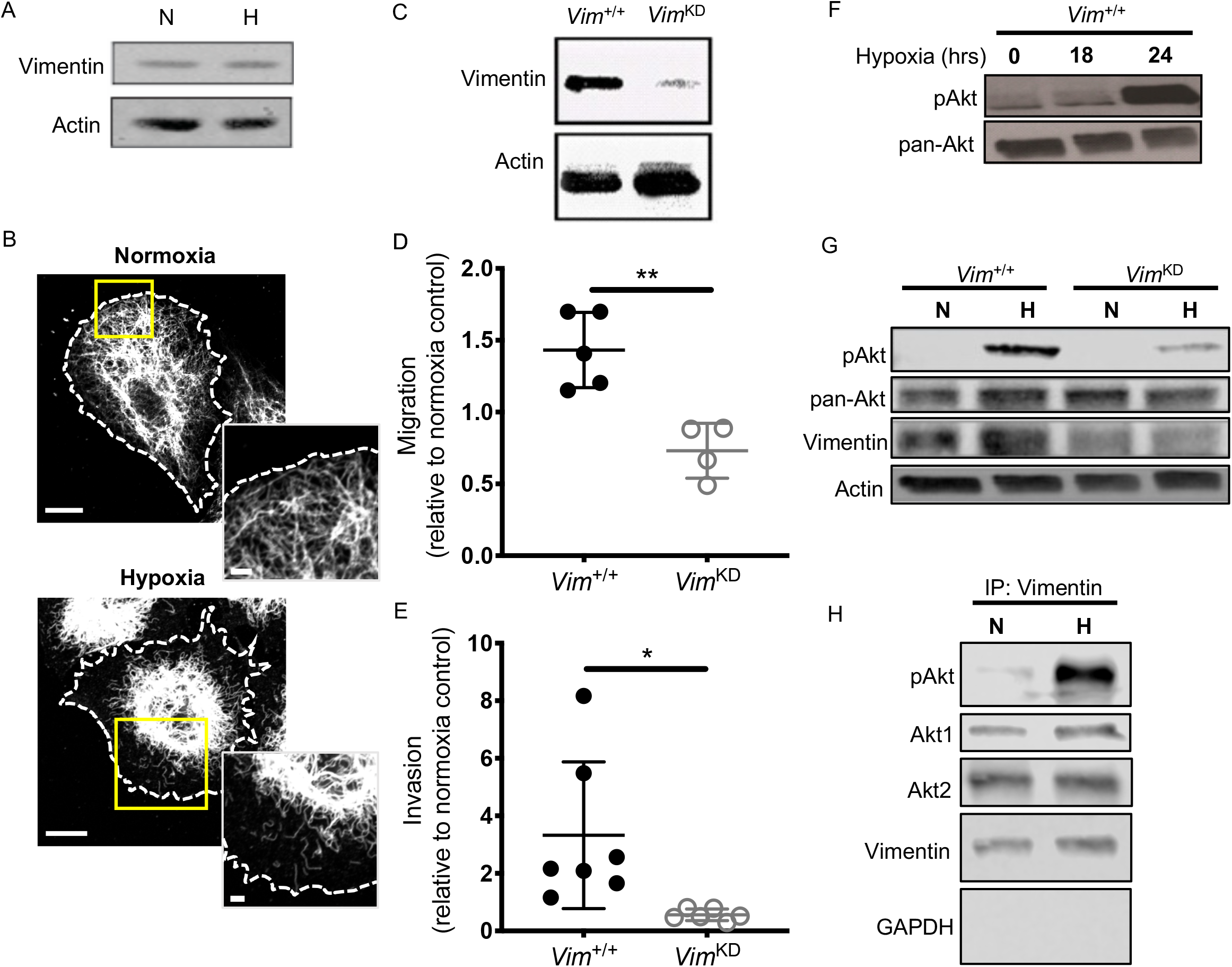
Vimentin is required for hypoxia-mediated cell migration and invasion. (**A**) Vimentin wild-type (*Vim*^+/+^) A549 cells were cultured in normoxic (N; 20% O_2_) or hypoxic (H; 1.5% O_2_) conditions for 24 hours. Total cell lysates were collected, separated by SDS-PAGE, and immunoblotted with antibodies against vimentin and actin. Representative confocal images of A549 cells exposed to normoxic or hypoxic conditions for 24 hours. Cells were fixed and stained for vimentin (white). A phase contrast image was used to identify cell borders (dashed line). Scale bars: 10 μm (*whole cells*), 2 μm *(inset*). (**C**) *Vim*^+/+^ A549 cells were treated with a retroviral vector expressing shRNA against vimentin (*Vim*KD). Total cell lysates were collected, separated by SDS-PAGE, and immunoblotted with antibodies against vimentin and actin. (**D, E**) Cells were cultured in hypoxic conditions, and migration over 6 hours (**D**, n=3–4) and invasion over 24 hours (**E**, n=6–8) were quantified. Each data point represents an independent experiment normalized to an average normoxic control. Data were compared using a two-tailed t-test and are presented as the mean ± standard deviation (*p<0.05; **p<0.01). (**F**) *Vim*^+/+^ cells were exposed to hypoxia for the indicated time; total cell lysates were collected, separated by SDS-PAGE, and immunoblotted with antibodies against phosphorylated Akt (pAkt; Ser473) and pan-Akt. (**G**) *Vim*^+/+^ and *Vim*^KD^ cells were exposed to hypoxia or normoxia for 24 hours. Total cell lysates were collected, separated by SDS-PAGE, and immunoblotted with antibodies against vimentin, pan-Akt, pAkt, and actin. (**H**) Vimentin was immunoprecipitated from total protein extracts derived from A549 cells exposed to normoxic or hypoxic conditions for 24 hours. Proteins were separated by SDS-PAGE and immunoblotted with antibodies against pAkt, Akt1, Akt2, vimentin, and GAPDH.

### Vimentin is required for lung cancer metastasis

The cell-autonomous ability of vimentin-expressing cells to metastasize *in vivo* was assessed using an allograft tumor model (**Figure 7A**). *Luc*-*KPV*^+/+^ cells, a luciferase expressing cell line that reproducibly colonizes to the lung following subcutaneous injection (44, 45), were transfected with CRISPR/Cas9 vimentin knockout plasmid to generate *Luc*-*KPV*^−/−^ cells (***SI Appendix***, **S5A**). Briefly, nude mice were subcutaneously injected in the right flank with either *Luc*-*KPV*^+/+^ or *Luc*-*KPV*^−/−^ cells (**Figure 7A**). Flank tumor volume and tumor radiance were measured weekly; there was no significant difference in either tumor volume or radiance between *Luc*-*KPV*^+/+^and *Luc*-*KPV*^−/−^ conditions during weeks 1–3. At week 3, flank tumors were excised (**Figure 7B**, ***SI Appendix***, **S4B**).

**Figure 7.**
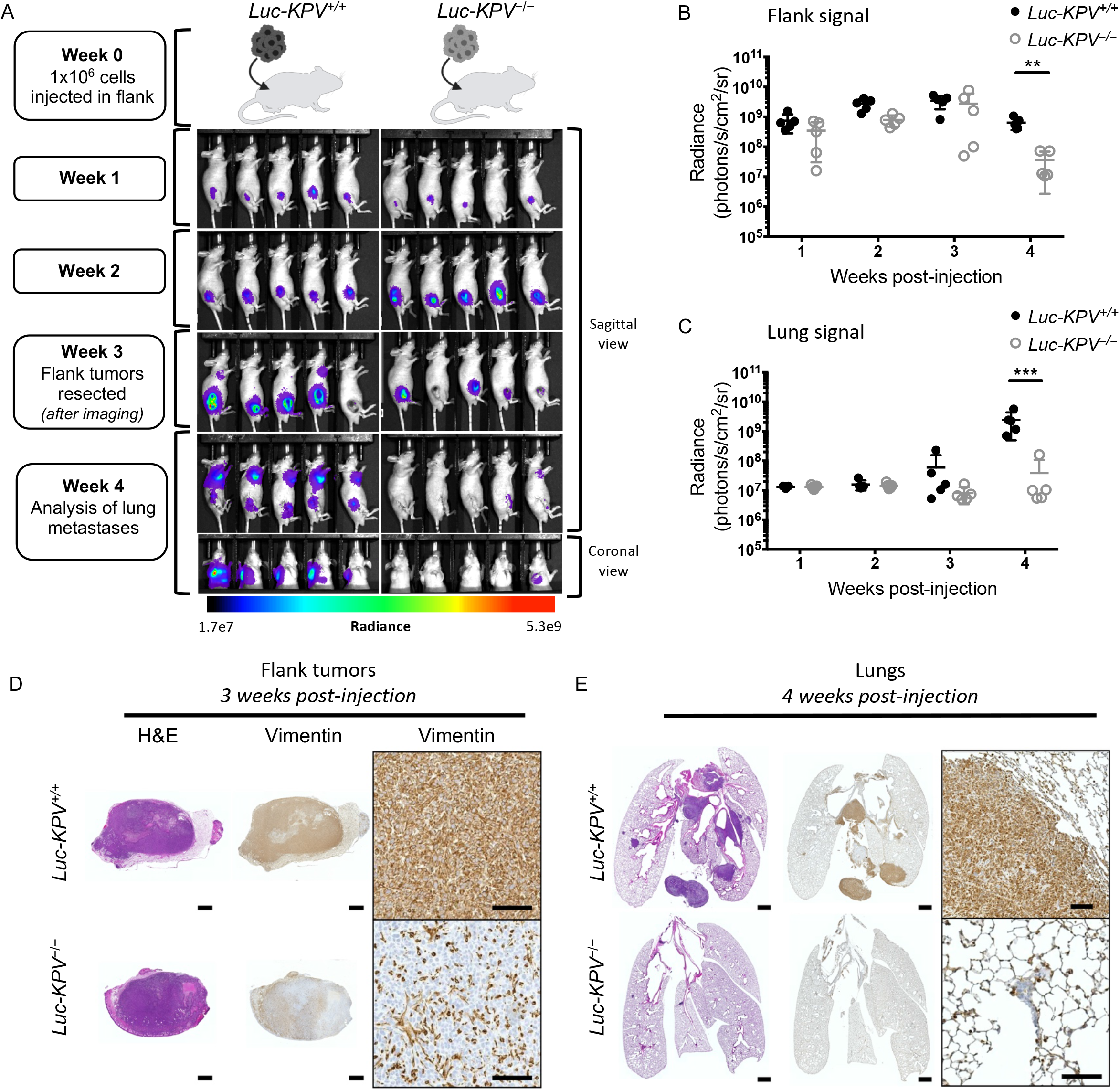
Vimentin is required for accelerated lung cancer metastasis. (**A**) Schematic of experimental design and accompanying IVIS images. A total of 1×106 *KPV*^+/+^ or *KPV*^−/−^ cells labeled with luciferase (*Luc-KPV*^+/+^ or *Luc-KPV*^−/−^, respectively) were injected subcutaneously into the right flank of nude mice. At 3 weeks post-injection, primary tumors were removed and lung metastases were tracked for an additional 1 week. Shown are representative IVIS images of mice (n=5 per group). For week 4, both sagittal and coronal views are shown. Coronal view was acquired after masking the flank tumor to minimize bleed-through of the signal. Intensity overlay shows the accumulation of luciferase-labeled cells. Luciferin signal was quantified from (**B**) primary flank tumors and (**C**) lungs. Lung radiance was quantified from masked images. Unpaired t-tests were used to compare *Luc-KPV*^+/+^ and *Luc-KPV*^−/−^ conditions at each time point (**p<0.01; ***p<0.001). Flank tumors (**D**) harvested at week 3 and lungs (**E**) harvested at week 4 were fixed, sectioned, and subjected to H&E staining and vimentin immunohistochemical staining. Positive vimentin staining is brown, and nuclei are blue. Scale bars: 1 mm (whole tumor/lung, *left*), 50 μm (inset, *right*).

At week 4 after injection, *Luc*-*KPV*^+/+^ cells colonized to the lung in 5 of 5 nude mice, whereas *KPV*^−/−^ cells failed to colonize to the lung in 4 of 5 nude mice (**Figure 7A-B**, ***SI Appendix***, **S5D-E**). Mice injected with *KPV*^+/+^ cells had considerable lung tumor burdens, as assessed by IVIS imaging and H&E staining (**Figure 7C and E**, ***SI Appendix***, **S5E**). In contrast, *KPV*^−/−^ cells, on average, failed to form lung tumors. When quantified, the metastatic signal in the lung was significantly higher in *KPV*^+/+^ mice (2.45E9 ± 1.95E9 photons•cm^−2^sr^−1^sec^−1^) than *KPV*^−/−^ mice (3.87E7 ± 6.90E7 photons•cm^−2^sr^−1^sec^−1^). Flank tumors that were removed at week 3 after injection were stained with H&E and immunostained for vimentin (**Figure 7D**). *KPV*^+/+^ cells formed dense tumors that displayed uniform vimentin expression. *KPV*^−/−^ cells also formed dense tumors; surprisingly, some cells within the tumor expressed vimentin. Based on their spindly or round shapes, we inferred that these cells were infiltrating fibroblasts or macrophages, two cell types which canonically express vimentin. The lungs of *KPV*^+/+^-injected mice displayed large, vimentin-positive metastatic lesions (**Figure 7E**). In contrast, the few metastatic tumors that formed in the lungs with *KPV*^−/−^ cells were sparse and small; as expected, these tumors did not express vimentin. Therefore, we conclude that vimentin is required for the rapid metastatic spread of murine lung adenocarcinoma cells. Furthermore, this effect is cell-autonomous. By injecting *KPV*^+/+^ and *KPV*^−/−^ cells into mice that have normal, vimentin-expressing stromal cells, we show that vimentin-expressing cells in the tumor microenvironment are not sufficient to promote the metastatic spread of vimentin-null cancer cells.

## Discussion

Clinically, vimentin expression correlates with increased metastatic potential (24), high nuclear grade (46), and poor overall survival across most solid tumor types including lung, prostate, and breast cancers (6–8). Vimentin has also been implicated in many aspects of cancer initiation and progression, including tumorigenesis, EMT, and the metastatic spread of cancer (9). Many of these reports relied on *in vitro* experiments comparing cultured cells derived from WT and *Vim^−/−^* mice. Our data provides causal evidence that vimentin is required for the metastasis of *Kras*-mutant, *Tp53*-null lung cancer cells *in vivo*. Data from the *KPV*^−/−^ GEMM show that vimentin is required for metastasis and tumor progression (**Figures 1 and 7**), as *KPV*^−/−^ mice had decreased lung tumor burden, lower grade tumors, and no metastasis from primary tumors in the flank to the lung (**Figures 1C-D and 7A-C**). Consistent with the decreased metastatic rates, we observed a survival advantage in the *KPV*^−/−^ mice (**Figure 1B**). These results were recapitulated in *KPV*^+/+^ mice by disrupting vimentin filaments with withaferin A treatment two weeks post tumor initiation (**Figure 4**). Collectively, these data provide evidence that vimentin is integral in the progression and metastasis of lung cancer.

The epithelial-to-mesenchymal transition (EMT) is the canonical mechanism by which cancer cells lose their epithelial morphology, form invadopodia and degrade the surrounding basement membrane to promote the invasive spread of cancer (9, 15). Several studies support the notion that vimentin functions as a positive regulator of EMT and that the upregulation of vimentin expression in epithelial cells is a prerequisite for EMT induction in malignant tumors (9, 13, 47–49). In this respect, it has been proposed that vimentin intermediate filaments provide a scaffold for the recruitment of transcription factors, such as Slug and Twist. Specifically, vimentin interacts with Slug to recruit ERK, which promotes the phosphorylation of Slug that is required for the initiation of the EMT (13). Similarly, when transforming growth factor β is used to activate the Smad-mediated EMT in primary alveolar epithelial cells, the shape changes characteristic of the EMT are directly associated with a rapid induction of vimentin expression regulated by a Smad-binding-element located in the 5′ promotor region of the *Vim* gene (14). Presently, we used RNA-seq to show that *KPV*^−/−^ cells derived from primary lung tumors display a distinct transcriptional phenotype, which is characterized by the suppression of genes directly involved in EMT, invasion and migration (**Figure 2**).

To invade into the surrounding tissue, an invasive tumor cell will first form invadopodia and degrade the surrounding basement membrane; vimentin is required for invadopodia formation (15). We showed that *KPV*^−/−^ cells fail to invade the surrounding extracellular matrix using a 3-dimensional experimental approach (**Figure 3B-C**). We previously reported that the transient expression of vimentin in epithelial cells, which typically express type I and type II keratin intermediate filaments causes epithelial cells to be transformed into mesenchymal cells, which is accompanied by changes in cell shape, increased cell motility and focal adhesion dynamics (11, 14). Direct evidence supporting the role of vimentin in the migration of *Kras*-mutant, *Tp53*-null lung cancer cells was demonstrated when vimentin expression was disrupted genetically (e.g. *KPV*^−/−^ cells and *Vim^KD^*) and pharmacologically (WFA) resulting in impaired migration. Importantly, *KPV*^−/−^ cells implanted in the flank of nude mice also failed to invade and migrate away from the primary tumor (**Figure 7**).

WFA and hypoxia treatment modulate cell motility of *Kras*-mutant, *Tp53*-null lung cancer cells in a vimentin-dependent manner. Cell motility decreases following WFA treatment and increases following hypoxia exposure, despite the seemingly similar effect WFA and hypoxia have on vimentin filament architecture (**Figure 3D and 6D**). These differences are likely due to vimentin phosphorylation, which regulates processes underlying cell motility in normal and cancer cells (50). WFA results in phosphorylation at Ser55/56 and hyperphosphorylation at Ser38/39 (51, 52). At Ser38/39, phosphorylation confers protection of the vimentin filament from caspase cleavage, while phosphorylation at Ser55/56 increases vimentin degradation which decreases the cell’s ability to invade and spread. On the other hand, hypoxia activates Akt, which binds to the head region of vimentin, resulting in the phosphorylation of vimentin at Ser38/39 (17). This interaction leads to hypoxia-mediated increases in cancer cell motility *in vitro*, as well as tumor and metastasis growth *in vivo*. Furthermore, Akt1 phosphorylation of vimentin offers a level of protection against caspase-mediated proteolysis, which allows for retention of mature vimentin filaments that can further contribute to cell motility (17).

We found that loss of vimentin alters the metabolic phenotype of *Kras*-mutant, *Tp53*-null lung cancer cells (**Figure 5**). Our data shows that *KPV*^−/−^ cells accumulate high levels of glutathione, which we hypothesize is due to an elevated response to oxidative stress. This is in line with a previous finding that vimentin-null cells produce higher levels of ROS compared to wildtype cells (53). Additionally, treatment with WFA induces ROS production in epithelial-derived cancer cells, suggesting that disruption of the vimentin intermediate filament network increases ROS generation (54). Furthermore, vimentin cooperates with ROS in production of collagen and cell alignment, functions that are necessary for directional cell motility (55). To that end, ROS-mediated vimentin reorganization, as shown in **Figure 6B**, allows for increased contraction, which could contribute to the force generation required for efficient migration and invasion (56). In cancers harboring a mutant *Kras* oncogene, mitochondrial production of ROS is required for tumor growth (57). Through a Rac1-mediated interaction with the mitochondria, phosphorylation of vimentin at Ser55/56 can mediate mitochondrial motility, leading to a decrease in mitochondrial membrane potential and vimentin-mediated protection of mitochondria from ROS (58–60). In vimentin-null cells, which have diminished mitochondrial function, a dysregulated oxidative stress response might account for the decreased tumor burden observed in *KPV*^−/−^ compared to *KPV*^+/+^ lungs. Additionally, we observed that genes associated with the OXPHOS hallmark gene set are negatively enriched in *KPV*^−/−^ cells compared to *KPV*^+/+^ cells (***SI Appendix***, **S3D**). We therefore reason that the possibly altered mitochondrial function in *KPV*^−/−^ cells is reflected in reduced OXPHOS. *KPV*^−/−^ may then rely on glycolysis for ATP production. Accordingly, we found that the glycolysis hallmark gene set was positively enriched in *KPV*^−/−^ cells (***SI Appendix***, **S3D**). Furthermore, in lung adenocarcinoma tissue, both oxidative phosphorylation (OXPHOS) and glycolysis are upregulated compared to normal adjacent lung tissue, suggesting that both pathways are associated with the disease (61). Together, these data suggest further exploration of the role of vimentin in cancer metabolism.

In the *Kras*-mutant, *Tp53*-null model of lung cancer, global vimentin depletion confers a survival advantage. Additionally, by treating *KPV*^+/+^ mice with WFA following tumor development, we show that the delayed tumor growth and metastasis observed in *KPV*^−/−^ mice in **Figure 1** is not due solely to delayed onset of tumor growth, but rather to attenuated growth kinetics in tumor cells that lack an intact vimentin network. Although vimentin-null mice were first reported to display no obvious phenotype, these data and others suggest that loss of vimentin is protective against a range of challenges including lung cancer, lipopolysaccharides, bleomycin, asbestos, bacterial meningitis, cerebral ischemia, and acute colitis (26, 38, 62–64). While this study adds to the body of knowledge on the phenotype of the vimentin-null mouse, the global knockout model presents limitations. Namely, in addition to lacking vimentin in cancer cells, *KPV*^−/−^ mice have vimentin-null stromal and immune compartments. Because vimentin is canonically expressed in mesenchymal, hematopoietic, and endothelial cells, which make up a large population of the tumor microenvironment (TME), loss of vimentin in these cells likely contributes to the decreased tumor burden in *KPV*^−/−^ mice compared to *KPV*^+/+^ mice (**Figure 1)**. To ensure that vimentin is sufficient in cancer cells to promote metastasis, we injected *KPV*^+/+^ or *KPV*^−/−^ cells into wildtype nude mice, which lack T cells but retain innate immune cells. We observed recruited mesenchymal and immune cells in subcutaneous flank tumors (**Figure 7D**), suggesting that the vimentin-positive TME in this model participates in growth of the primary tumor. Despite being in the presence of vimentin-expressing stromal and immune cells, *KPV*^−/−^ cells failed to metastasize. Therefore, while other groups have found that vimentin deficiency impairs function in cancer-associated fibroblasts and immune cells such as macrophages and T cells (49, 62, 65, 66), a vimentin-expressing microenvironment is not sufficient to promote metastasis in the time frame evaluated. However, more research is needed to fully understand how vimentin in non-cancer cells may be synergistically controlling metastasis. Our group has previously shown that vimentin is required for the production of mature interleukin 1β (IL-1β), a mediator of cancer growth and metastasis (62, 67). The cytokine IL-1β further promotes tumor-associated macrophage infiltration, which could explain the decreased recruitment of CD45+ cells in *KPV*^−/−^ tumors compared to KPV^+/+^ tumors (***SI Appendix***, **S2C**) (68). To better understand how vimentin participates in different compartments of the TME, we recognize that animal models with immune-, mesenchymal-, and epithelial-specific deletion of vimentin will need to be created.

This work gives physiological context to a large range of clinical data that links vimentin to cancer progression (6, 24, 69). There is also a wealth of *in vitro* research that provide a number of vimentin-dependent mechanisms related to cancer metastasis. Broadly, these mechanisms include interacting with actin to form lamellipodia and invadopodia, stabilization of collagen mRNA, guiding microtubules to control cell polarity, and aligning actin-potentiated traction forces (15, 16, 70–73). Ultimately, our findings provide *in vivo* context for a multitude of clinical and *in vitro* reports by showing that vimentin is required for lung cancer metastasis. Through genetic and chemical interference, we have identified vimentin as a potential clinical target for metastatic lung cancer.

## Materials and Methods

### Murine lung cancer model

All animal experiments were approved by Northwestern University’s Institutional Animal Care and Use Committee (IACUC). Sex-matched 6–10-week-old mice were used for all *in vivo* experiments. *LSL*-*Kras*^G12D/+^;*Tp53*^flox/flox^ (*KPV*^+/+^) mice were bred as described by DuPage and colleagues and were generously gifted to us by Dr. Navdeep Chandel (Northwestern University, Chicago, IL) (25). Vimentin-knockout mice were a gift from Albee Messing (University of Wisconsin, Madison, WI). Vimentin-knockout mice were crossed with *KPV*^+/+^ mice to create *KPV*^−/−^ mice. *KPV*^+/+^, *KPV*^−/−^, and the validated *Rosa26-LSL-LacZ* mice were administered adenovirus expressing Cre recombinase (Ad-Cre; ViraQuest) or a null adenovirus (Ad-Null) via intratracheal instillation (1 × 10^9^ pfu unless otherwise noted) under isoflurane anesthesia (28). Survival was monitored daily. Weight was monitored weekly.

### Magnetic resonance imaging

Scheduled magnetic resonance imaging (MRI) was performed at Northwestern University Center for Translational Imaging (Chicago, IL) via a 7-tesla system (Clinscan, Bruker) using a four-channel mouse body coil at set time points (2, 6, and 10 weeks after Ad-Cre administration). In order to permit tolerance to imaging, the mice were anesthetized with isoflurane (2% isoflurane in oxygen for induction, followed by 1.5–2% via nose cone for maintenance during imaging). Pulse oximetry and respiration were recorded and used to trigger the MRI in order to avoid motion artifacts. Turbo Multi Spin Echo imaging sequence was used in conjunction with respiratory triggering to acquire *in vivo* MRI coronal images covering all the lung area and portions of abdomen, including liver and kidneys (ST = 0.5 mm, In plane = 120 μm, TR =1000 msec, TE= 20 msec). Gradient Echo sequence was used with cardiac triggering (using pulse oximeter rate) covering the lung area transversally (ST = 0.5 mm, In plane = 120 μm, TR ~20 msec, TE ~ 2 msec). Jim software was used to quantify tumor burden (Xinapse).

### Immunohistochemistry

Mice were anesthetized and lungs were perfused via the right ventricle with 4% paraformaldehyde in phosphate-buffered saline (PBS). A 20-gauge angiocatheter was sutured into the trachea, heart and lungs were removed en bloc, and then lungs were inflated with 0.8 mL of 4% paraformaldehyde at a pressure not exceeding 16 cm H_2_O. Tissue was fixed in 4% paraformaldehyde in PBS overnight at 4°C, then processed, embedded in paraffin, and sectioned (4–5 μm). Tissue sections were stained with hematoxylin and eosin (H&E) or used for immunohistochemistry. After rehydration, tissues were subjected to antigen retrieval in 10 mM sodium citrate (pH = 6.0) with 0.05% Tween-20 for 20 minutes at 96–98°C, followed by 20 minutes of cooling. Tissue sections were blocked in 3% hydrogen peroxide for 5 min, then a Vector Laboratories avidin/biotin blocking kit (SP-2001), Vectastain ABC kit (PK-4001), and 3,3′-diaminobenzidine (DAB) peroxidase substrate kit (SK-4100) were used according to the manufacturer’s protocols. Nuclei were counterstained with hematoxylin (Thermo Scientific 72604) and treated with bluing solution (Thermo Scientific 7301), and then coverslips were mounted with Cytoseal 60 (Thermo Scientific 8310-4). A TissueGnostics automated slide imaging system was used to acquire whole-tissue images and measure area.

### Cell isolation and culture

*KPV*^+/+^ and *KPV*^−/−^ mice were treated with Ad-Cre as described above; after 6 weeks, mice were sacrificed and lung tumors were excised. Tissue was dissociated into a single cell suspension in 0.2 mg/mL DNase and 2 mg/mL collagenase D and was filtered through a 40 μm filter. Cells then underwent two rounds of selection. First, cells were treated with anti-CD45 magnetic beads (Miltenyi Biotec, 130-052-301) and were passed through a magnetic column. CD45-negative cells were then subjected to anti-EPCAM magnetic beads (Miltenyi Biotec, 130-105-958) and underwent positive selection. CD45-negative, EPCAM-positive cells were expanded *in vitro* and were used in experiments between passages 1 and 10. Cells derived from a human lung adenocarcinoma (A549, CCL-185) were obtained from the American Type Culture Collection (ATCC, Manassas, VA). All cells were maintained in Dulbecco’s modified Eagle medium (DMEM) supplemented with 10% fetal bovine serum, 100 U/mL penicillin, 100 μg/mL streptomycin, and HEPES buffer. All cells were grown in a humidified incubator of 5% CO_2_/95% air at 37°C (unless otherwise noted).

### Polymerase chain reaction

Mice were infected with Ad-Null or Ad-Cre; at 2, 8, and 12 wpi, mice were sacrificed and lungs were harvested. Lungs were lysed, and DNA was extracted and amplified by polymerase chain reaction (PCR) using the following primers: *Kras* forward, GGC CTG CTG AAA ATG ACT GAG TAT A; *Kras* reverse, CTG TAT CGT CAA GGC GCT CTT; *Kras-G12D* forward, CTTGTGGTGGTTGGAGCTGA; and *Kras-G12D* reverse, TCCAAGAGACAGGTTTCTCCA. DNA products were run on an agarose gel and imaged with the Li-Cor Odyssey imaging system.

### Western blotting

Western blot analysis was utilized to quantify protein levels in cell lysates. The protein was separated using 12% sodium dodecyl sulfate polyacrylamide gel electrophoresis (SDS-PAGE) and transferred onto nitrocellulose membranes. Membranes were then blocked with Odyssey blocking buffer (Li-Cor Biosciences) and subsequently incubated with the appropriate primary antibodies overnight at 4°C. IRDye secondary antibodies were then used (Li-Cor Biosciences, 1:10,000) for 2 hours at room temperature. Images of blots were acquired using the Li-Cor Odyssey Fc Imaging System.

### RNA-sequencing

Tumor cells were isolated from *KPV*^+/+^ and *KPV*^−/−^ mice at 6 wpi and underwent CD45-negative, EpCAM-positive magnetic-activated cell sorting (MACS) selection as described above. Cells were cultured for one passage, then lysed using RLT lysis buffer (Qiagen), and total RNA was isolated with the RNeasy Plus Mini Kit (Qiagen). Quality of RNA was confirmed with a TapeStation 4200 (Agilent); all samples had an RNA integrity number (RIN) score equal to or greater than 9.8. Next, mRNA was isolated via poly(A) enrichment (NEBNext). Libraries were prepared using NEBNext RNA Ultra chemistry (New England Biolabs). Sequencing was performed on an Illumina NextSeq 500 using a 75-cycle single-end high-output sequencing kit. Reads were demultiplexed (bcl2fastq), and fastq files were aligned to the mm10 mouse reference genome with TopHat2. Htseq was used to obtain counts. The resulting data were filtered, and differentially expressed genes (DEGs) were identified using the edgeR package. DEGs were selected using a false discovery rate (FDR) cutoff of <0.05, with a 1.0-fold change cutoff for pairwise comparison. K-means clustering and heat map visualization was performed using the Morpheus web tool (https://software.broadinstitute.org/morpheus). Enrichment analysis was performed using Gorilla (74, 75).

### Withaferin A treatments

Withaferin A (WFA) was purchased from Enzo Life Sciences and dissolved in dimethyl sulfoxide (DMSO; Sigma-Aldrich) to a final concentration of 5 μM unless noted otherwise. For *in vivo* experiments, jelly pellets were utilized to provide an oral, voluntary method of drug delivery. Using a 24-well flat-bottom tissue culture plate as the jelly mold, WFA (4 mg/kg in DMSO) or vehicle control (DMSO only) were combined with gelatin and Splenda for flavoring as described elsewhere (76). Tumor development was induced with Ad-Cre as described above. Two weeks following Ad-Cre administration, mice were fed jelly pellets every other day for 4 weeks. Survival was tracked daily and weight was measured weekly.

### Scratch wound assay

Cells were grown to 100% confluence in 6-well plates. A pipette tip was used to make a single scratch in the monolayer. The cells were washed with 1× PBS to remove debris and imaged at 0 hours and 6 hours. For WFA conditions, WFA or DMSO was added at 0 hours (when the scratch was created). Rate of cell migration was calculated using ImageJ software. Results were normalized to the initial wound area at 0 hours.

### Matrigel invasion assay

Transwell inserts with 8 μm pores were coated with Matrigel (200 μg/mL), and 5 × 10^4^ *KPV*^+/+^ or *KPV*^−/−^ cells were seeded atop each transwell in serum-free media. For all experiments with A549 cells, the cells were serum-starved for 24 hours and were then plated at a concentration of 1 × 10^5^ cells per transwell. For WFA experiments, cells were resuspended in WFA or DMSO containing media directly before being seeded in transwells. Media containing 10% fetal bovine serum was added to the bottom well to serve as a chemoattractant. Cells were placed at 37°C for 48 hours (24 hours for A549 cells). Following incubation, the Matrigel with the cells remaining on the upper surface of the transwell was removed with a cotton swab. The cells remaining on the bottom of the membrane were fixed in 2% paraformaldehyde and incubated with Hoechst nuclear dye (Invitrogen; 1:10,000 in 1× PBS). Five random 10× magnification fields were imaged, and the average number of cells per field was quantified; this average is reported as the “invasive index.”

### Spheroid culture

Spheroids were generated as described by Gilbert-Ross et al. (77). Briefly, cells were grown in Nunclon Sphera 96-well plates (Thermo-Fisher Scientific) at a concentration of 3000 cells per well. After 3 days in culture, cells were transferred using a wide-bore pipette tip to 2 mg/mL collagen (Corning) in 4-well LabTek plates (Nunc). Collagen was allowed to gel at 37°C for 1 hour; then, complete media was added to the spheroids. Gels were imaged using a Ti2 widefield microscope (Nikon) at 0, 24, and 48 hours. Spheroid area was quantified using Fiji software. Reported spheroid area values are normalized to 0-hour spheroid area of the same spheroid.

### Hypoxia conditions

A hypoxic environment was created *in vitro* by culturing cells in 1.5% O_2_, 93.5% N_2_, and 5% CO_2_ in a humidified variable aerobic workstation (InVivo O2; BioTrace International, Muncie, IN). Before experimentation, cell culture medium was allowed to equilibrate to oxygen levels overnight.

### Metabolomics

*KPV*^+/+^ and *KPV*^−/−^ cells were grown in 6-well plates. High-performance liquid chromatography (HPLC) grade methanol (80% in water) was added to cells, and plates were incubated at −80°C for 20 minutes. Lysates were collected and centrifuged, and the supernatant was collected and analyzed by High-Performance Liquid Chromatography and High-Resolution Mass Spectrometry and Tandem Mass Spectrometry (HPLC-MS/MS). Specifically, system consisted of a Thermo Q-Exactive in line with an electrospray source and an Ultimate3000 (Thermo) series HPLC consisting of a binary pump, degasser, and auto-sampler outfitted with a Xbridge Amide column (Waters; dimensions of 4.6 mm × 100 mm and a 3.5 μm particle size). The mobile phase A contained 95% (vol/vol) water, 5% (vol/vol) acetonitrile, 20 mM ammonium hydroxide, 20 mM ammonium acetate, pH = 9.0; B was 100% Acetonitrile. The gradient was as following: 0 min, 15% A; 2.5 min, 30% A; 7 min, 43% A; 16 min, 62% A; 16.1-18 min, 75% A; 18-25 min, 15% A with a flow rate of 400 μL/min. The capillary of the ESI source was set to 275 °C, with sheath gas at 45 arbitrary units, auxiliary gas at 5 arbitrary units and the spray voltage at 4.0 kV. In positive/negative polarity switching mode, an *m*/*z* scan range from 70 to 850 was chosen and MS1 data was collected at a resolution of 70,000. The automatic gain control (AGC) target was set at 1 × 10^6^ and the maximum injection time was 200 ms. The top 5 precursor ions were subsequently fragmented, in a data-dependent manner, using the higher energy collisional dissociation (HCD) cell set to 30% normalized collision energy in MS2 at a resolution power of 17,500. Besides matching m/z, metabolites are identified by matching either retention time with analytical standards and/or MS2 fragmentation pattern. Data acquisition and analysis were carried out by Xcalibur 4.1 software and Tracefinder 4.1 software, respectively (both from Thermo Fisher Scientific). For each sample, peak area of each metabolite was normalized to total ion count per sample. Data were log-transformed and compared with a two-tailed, unpaired t-test. Data was analyzed with MetaboAnalyst software (78).

### Preparation of cells for subcutaneous flank injection

*KPV*^+/+^ cells labeled with luciferase (*Luc*-*KPV*^+/+^ cells) were a generous gift from Dr. Navdeep Chandel. To create *Luc*-*KPV*^−/−^ cells, *Luc*-*KPV*^+/+^ cells were transfected with a commercially available CRISPR/Cas9 vimentin knockout plasmid according to manufacturer’s directions (Santa Cruz Biotechnology sc-423676).

### Tracking of tumor growth in subcutaneous flank injection model

Male nude (NU/J) mice were purchased from Jackson Laboratories; 8–12-week-old mice were anesthetized with 2% isoflurane in oxygen and were given a subcutaneous injection of cells (1 × 10^6^ cells in 100 μL of 1× PBS) on their right flanks. Weight and tumor volume were monitored weekly. For IVIS imaging, mice were injected with 150 mg of D-luciferin per kilogram of body weight (PerkinElmer 770504). After 10 minutes, IVIS images were captured. At week 3 post-injection, tumors were removed. Briefly, mice were anesthetized with ketamine (100 mg/kg body weight) and xylazine (10 mg/kg body weight). Tumor area was disinfected with 70% ethanol and iodide solution. Tumors were excised and placed in 4% paraformaldehyde for immunohistochemistry. Wounds were closed with simple interrupted nylon sutures (Ethilon). Mice were monitored until they recovered from anesthesia; they were then housed singly and treated with Meloxicam as an analgesic. The following week, mice underwent a final IVIS imaging session and were then sacrificed.

### Immunofluorescence confocal microscopy

For all immunofluorescent immunocytochemistry experiments, cells were grown on no. 1 glass coverslips. Following treatment, *KPV*^+/+^ and *KPV*^−/−^ cells were fixed in methanol for 3–5 minutes. A549 cells were fixed with 2% paraformaldehyde for 7–10 minutes. *KPV*^+/+^ and *KPV*^−/−^ cells were blocked in 5% normal goat serum (NGS) for 1 hour at room temperature. A549 cells were blocked with 1.5% NGS for 30 minutes at 37°C. Cells were then treated with the indicated primary antibodies overnight at 4°C. Cells were washed twice in PBS with 0.10% Tween-20 for 3 minutes each and treated with secondary antibodies conjugated with Alexa Fluor 488 (Invitrogen A-11039, 1:200) and/or Alexa Fluor 568 (Invitrogen A-11004, 1:200), as well as Hoechst nuclear dye (Invitrogen H3570, 1:10,000). Coverslips were mounted and sealed. A Nikon A1R+ laser scanning confocal microscope equipped with a 60× and 100× objective lens was used to acquire images. For experiments with A549 cells, a Zeiss LSM 510 laser scanning confocal microscope equipped with a 63× objective lens was used to acquire images. Nikon NIS-Elements software and ImageJ were used for image processing.

### Reagents

All antibodies used are summarized in **Supplemental Table 1**.

## Acknowledgments

Histology services were provided by the Northwestern University Mouse Histology and Phenotyping Laboratory which is supported by NCI P30-CA060553 awarded to the Robert H Lurie Comprehensive Cancer Center. Imaging work was performed at the Northwestern University Center for Advanced Microscopy generously supported by NCI CCSG P30 CA060553 awarded to the Robert H Lurie Comprehensive Cancer Center. Metabolomics services were performed by the Metabolomics Core Facility at Robert H. Lurie Comprehensive Cancer Center of Northwestern University. We would like to thank Hiam Abdala-Valencia for performing RNA-sequencing. All graphic design was created with BioRender.com.

## Notes

Supported by: A.B. and K.W. were supported by NIH/NHLBI T32 HL076139; K.R.A. was supported by David and Christine Cugell Fellowship; K.M.R. was supported by NIH P01HL071643, R01HL128194, P01AG049665.

### Competing Interest Statement

The authors have declared no competing interest.

## References

1. Siegel RL, Miller KD, Jemal A. Cancer statistics, 2019. CA Cancer J Clin. 2019;69(1):7–34. Epub 2019/01/09. doi: 10.3322/caac.21551. PubMed PMID: 30620402.

2. Shajani-Yi Z, de Abreu FB, Peterson JD, Tsongalis GJ. Frequency of Somatic TP53 Mutations in Combination with Known Pathogenic Mutations in Colon Adenocarcinoma, Non-Small Cell Lung Carcinoma, and Gliomas as Identified by Next-Generation Sequencing. Neoplasia. 2018;20(3):256–62. Epub 2018/02/18. doi: 10.1016/j.neo.2017.12.005. PubMed PMID: 29454261; PMCID: PMC5849803.

3. Scoccianti C, Vesin A, Martel G, Olivier M, Brambilla E, Timsit JF, Tavecchio L, Brambilla C, Field JK, Hainaut P, European Early Lung Cancer C. Prognostic value of TP53, KRAS and EGFR mutations in nonsmall cell lung cancer: the EUELC cohort. Eur Respir J. 2012;40(1):177–84. Epub 2012/01/24. doi: 10.1183/09031936.00097311. PubMed PMID: 22267755.

4. Cox AD, Fesik SW, Kimmelman AC, Luo J, Der CJ. Drugging the undruggable RAS: Mission possible? Nat Rev Drug Discov. 2014;13(11):828–51. Epub 2014/10/18. doi: 10.1038/nrd4389. PubMed PMID: 25323927; PMCID: PMC4355017.

5. Cancer Genome Atlas Research N. Comprehensive molecular profiling of lung adenocarcinoma. Nature. 2014;511(7511):543–50. Epub 2014/08/01. doi: 10.1038/nature13385. PubMed PMID: 25079552; PMCID: PMC4231481.

6. Dauphin M, Barbe C, Lemaire S, Nawrocki-Raby B, Lagonotte E, Delepine G, Birembaut P, Gilles C, Polette M. Vimentin expression predicts the occurrence of metastases in non small cell lung carcinomas. Lung Cancer. 2013;81(1):117–22. Epub 2013/04/09. doi: 10.1016/j.lungcan.2013.03.011. PubMed PMID: 23562674.

7. Burch TC, Watson MT, Nyalwidhe JO. Variable metastatic potentials correlate with differential plectin and vimentin expression in syngeneic androgen independent prostate cancer cells. PLoS One. 2013;8(5):e65005. doi: 10.1371/journal.pone.0065005. PubMed PMID: 23717685; PMCID: PMC3661497.

8. Domagala W, Lasota J, Dukowicz A, Markiewski M, Striker G, Weber K, Osborn M. Vimentin expression appears to be associated with poor prognosis in node-negative ductal NOS breast carcinomas. Am J Pathol. 1990;137(6):1299–304. PubMed PMID: 1701960; PMCID: PMC1877729.

9. Kidd ME, Shumaker DK, Ridge KM. The role of vimentin intermediate filaments in the progression of lung cancer. Am J Respir Cell Mol Biol. 2014;50(1):1–6. Epub 2013/08/29. doi: 10.1165/rcmb.2013-0314TR. PubMed PMID: 23980547; PMCID: PMC3930939.

10. Dongre A, Weinberg RA. New insights into the mechanisms of epithelial-mesenchymal transition and implications for cancer. Nat Rev Mol Cell Biol. 2019;20(2):69–84. Epub 2018/11/22. doi: 10.1038/s41580-018-0080-4. PubMed PMID: 30459476.

11. Mendez MG, Kojima S, Goldman RD. Vimentin induces changes in cell shape, motility, and adhesion during the epithelial to mesenchymal transition. FASEB J. 2010;24(6):1838–51. Epub 2010/01/26. doi: 10.1096/fj.09-151639. PubMed PMID: 20097873; PMCID: PMC2874471.

12. Meng J, Chen S, Han JX, Qian B, Wang XR, Zhong WL, Qin Y, Zhang H, Gao WF, Lei YY, Yang W, Yang L, Zhang C, Liu HJ, Liu YR, Zhou HG, Sun T, Yang C. Twist1 Regulates Vimentin through Cul2 Circular RNA to Promote EMT in Hepatocellular Carcinoma. Cancer Res. 2018;78(15):4150–62. Epub 2018/05/31. doi: 10.1158/0008-5472.CAN-17-3009. PubMed PMID: 29844124.

13. Virtakoivu R, Mai A, Mattila E, De Franceschi N, Imanishi SY, Corthals G, Kaukonen R, Saari M, Cheng F, Torvaldson E, Kosma VM, Mannermaa A, Muharram G, Gilles C, Eriksson J, Soini Y, Lorens JB, Ivaska J. Vimentin-ERK Signaling Uncouples Slug Gene Regulatory Function. Cancer Res. 2015;75(11):2349–62. Epub 2015/04/10. doi: 10.1158/0008-5472.CAN-14-2842. PubMed PMID: 25855378.

14. Rogel MR, Soni PN, Troken JR, Sitikov A, Trejo HE, Ridge KM. Vimentin is sufficient and required for wound repair and remodeling in alveolar epithelial cells. FASEB J. 2011;25(11):3873–83. doi: 10.1096/fj.10-170795. PubMed PMID: 21803859; PMCID: PMC3205840.

15. Schoumacher M, Goldman RD, Louvard D, Vignjevic DM. Actin, microtubules, and vimentin intermediate filaments cooperate for elongation of invadopodia. J Cell Biol. 2010;189(3):541–56. Epub 2010/04/28. doi: 10.1083/jcb.200909113. PubMed PMID: 20421424; PMCID: PMC2867303.

16. Helfand BT, Mendez MG, Murthy SN, Shumaker DK, Grin B, Mahammad S, Aebi U, Wedig T, Wu YI, Hahn KM, Inagaki M, Herrmann H, Goldman RD. Vimentin organization modulates the formation of lamellipodia. Mol Biol Cell. 2011;22(8):1274–89. Epub 2011/02/25. doi: 10.1091/mbc.E10-08-0699. PubMed PMID: 21346197; PMCID: PMC3078081.

17. Zhu QS, Rosenblatt K, Huang KL, Lahat G, Brobey R, Bolshakov S, Nguyen T, Ding Z, Belousov R, Bill K, Luo X, Lazar A, Dicker A, Mills GB, Hung MC, Lev D. Vimentin is a novel AKT1 target mediating motility and invasion. Oncogene. 2011;30(4):457–70. Epub 2010/09/22. doi: 10.1038/onc.2010.421. PubMed PMID: 20856200; PMCID: PMC3010301.

18. Zelenko Z, Gallagher EJ, Tobin-Hess A, Belardi V, Rostoker R, Blank J, Dina Y, LeRoith D. Silencing vimentin expression decreases pulmonary metastases in a pre-diabetic mouse model of mammary tumor progression. Oncogene. 2017;36(10):1394–403. Epub 2016/08/30. doi: 10.1038/onc.2016.305. PubMed PMID: 27568979; PMCID: PMC5332535.

19. Hendrix MJ, Seftor EA, Seftor RE, Trevor KT. Experimental co-expression of vimentin and keratin intermediate filaments in human breast cancer cells results in phenotypic interconversion and increased invasive behavior. Am J Pathol. 1997;150(2):483–95. Epub 1997/02/01. PubMed PMID: 9033265; PMCID: PMC1858294.

20. Gilles C, Polette M, Zahm JM, Tournier JM, Volders L, Foidart JM, Birembaut P. Vimentin contributes to human mammary epithelial cell migration. J Cell Sci. 1999;112 (Pt 24):4615–25. Epub 1999/11/27. PubMed PMID: 10574710.

21. Messica Y, Laser-Azogui A, Volberg T, Elisha Y, Lysakovskaia K, Eils R, Gladilin E, Geiger B, Beck R. The role of Vimentin in Regulating Cell Invasive Migration in Dense Cultures of Breast Carcinoma Cells. Nano Lett. 2017;17(11):6941–8. doi: 10.1021/acs.nanolett.7b03358. PubMed PMID: 29022351.

22. Wang W, Yi M, Zhang R, Li J, Chen S, Cai J, Zeng Z, Li X, Xiong W, Wang L, Li G, Xiang B. Vimentin is a crucial target for anti-metastasis therapy of nasopharyngeal carcinoma. Mol Cell Biochem. 2018;438(1-2):47–57. Epub 2017/07/27. doi: 10.1007/s11010-017-3112-z. PubMed PMID: 28744809.

23. Chan SH, Tsai JP, Shen CJ, Liao YH, Chen BK. Oleate-induced PTX3 promotes head and neck squamous cell carcinoma metastasis through the up-regulation of vimentin. Oncotarget. 2017;8(25):41364–78. Epub 2017/05/11. doi: 10.18632/oncotarget.17326. PubMed PMID: 28489600; PMCID: PMC5522334.

24. Liu S, Liu L, Ye W, Ye D, Wang T, Guo W, Liao Y, Xu D, Song H, Zhang L, Zhu H, Deng J, Zhang Z. High Vimentin Expression Associated with Lymph Node Metastasis and Predicated a Poor Prognosis in Oral Squamous Cell Carcinoma. Scientific reports. 2016;6:38834. doi: 10.1038/srep38834. PubMed PMID: 27966589; PMCID: PMC5155220.

25. DuPage M, Dooley AL, Jacks T. Conditional mouse lung cancer models using adenoviral or lentiviral delivery of Cre recombinase. Nat Protoc. 2009;4(7):1064–72. Epub 2009/06/30. doi: 10.1038/nprot.2009.95. PubMed PMID: 19561589; PMCID: PMC2757265.

26. Colucci-Guyon E, Portier MM, Dunia I, Paulin D, Pournin S, Babinet C. Mice lacking vimentin develop and reproduce without an obvious phenotype. Cell. 1994;79(4):679–94. Epub 1994/11/18. doi: 10.1016/0092-8674(94)90553-3. PubMed PMID: 7954832.

27. Jackson EL, Olive KP, Tuveson DA, Bronson R, Crowley D, Brown M, Jacks T. The differential effects of mutant p53 alleles on advanced murine lung cancer. Cancer Res. 2005;65(22):10280–8. Epub 2005/11/17. doi: 10.1158/0008-5472.CAN-05-2193. PubMed PMID: 16288016.

28. Soriano P. Generalized lacZ expression with the ROSA26 Cre reporter strain. Nat Genet. 1999;21(1):70–1. Epub 1999/01/23. doi: 10.1038/5007. PubMed PMID: 9916792.

29. Whithaus K, Fukuoka J, Prihoda TJ, Jagirdar J. Evaluation of napsin A, cytokeratin 5/6, p63, and thyroid transcription factor 1 in adenocarcinoma versus squamous cell carcinoma of the lung. Arch Pathol Lab Med. 2012;136(2):155–62. Epub 2012/02/01. doi: 10.5858/arpa.2011-0232-OA. PubMed PMID: 22288962.

30. Simanshu DK, Nissley DV, McCormick F. RAS Proteins and Their Regulators in Human Disease. Cell. 2017;170(1):17–33. Epub 2017/07/01. doi: 10.1016/j.cell.2017.06.009. PubMed PMID: 28666118; PMCID: PMC5555610.

31. Nagalingam A, Kuppusamy P, Singh SV, Sharma D, Saxena NK. Mechanistic elucidation of the antitumor properties of withaferin a in breast cancer. Cancer Res. 2014;74(9):2617–29. Epub 2014/04/16. doi: 10.1158/0008-5472.CAN-13-2081. PubMed PMID: 24732433; PMCID: PMC4009451.

32. Suman S, Das TP, Moselhy J, Pal D, Kolluru V, Alatassi H, Ankem MK, Damodaran C. Oral administration of withaferin A inhibits carcinogenesis of prostate in TRAMP model. Oncotarget. 2016;7(33):53751–61. Epub 2016/07/23. doi: 10.18632/oncotarget.10733. PubMed PMID: 27447565; PMCID: PMC5288218.

33. Kakar SS, Parte S, Carter K, Joshua IG, Worth C, Rameshwar P, Ratajczak MZ. Withaferin A (WFA) inhibits tumor growth and metastasis by targeting ovarian cancer stem cells. Oncotarget. 2017;8(43):74494–505. Epub 2017/11/02. doi: 10.18632/oncotarget.20170. PubMed PMID: 29088802; PMCID: PMC5650357.

34. Bargagna-Mohan P, Hamza A, Kim YE, Khuan Abby Ho Y, Mor-Vaknin N, Wendschlag N, Liu J, Evans RM, Markovitz DM, Zhan CG, Kim KB, Mohan R. The tumor inhibitor and antiangiogenic agent withaferin A targets the intermediate filament protein vimentin. Chem Biol. 2007;14(6):623–34. Epub 2007/06/23. doi: 10.1016/j.chembiol.2007.04.010. PubMed PMID: 17584610; PMCID: PMC3228641.

35. Bollong MJ, Pietila M, Pearson AD, Sarkar TR, Ahmad I, Soundararajan R, Lyssiotis CA, Mani SA, Schultz PG, Lairson LL. A vimentin binding small molecule leads to mitotic disruption in mesenchymal cancers. Proceedings of the National Academy of Sciences of the United States of America. 2017;114(46):E9903–E12. Epub 2017/11/01. doi: 10.1073/pnas.1716009114. PubMed PMID: 29087350; PMCID: PMC5699095.

36. Mahesh PP, Retnakumar RJ, Mundayoor S. Downregulation of vimentin in macrophages infected with live Mycobacterium tuberculosis is mediated by Reactive Oxygen Species. Sci Rep. 2016;6:21526. Epub 2016/02/16. doi: 10.1038/srep21526. PubMed PMID: 26876331; PMCID: PMC4753491.

37. Tolstonog GV, Belichenko-Weitzmann IV, Lu JP, Hartig R, Shoeman RL, Traub U, Traub P. Spontaneously immortalized mouse embryo fibroblasts: growth behavior of wild-type and vimentin-deficient cells in relation to mitochondrial structure and activity. DNA Cell Biol. 2005;24(11):680–709. Epub 2005/11/09. doi: 10.1089/dna.2005.24.680. PubMed PMID: 16274292.

38. Mor-Vaknin N, Legendre M, Yu Y, Serezani CH, Garg SK, Jatzek A, Swanson MD, Gonzalez-Hernandez MJ, Teitz-Tennenbaum S, Punturieri A, Engleberg NC, Banerjee R, Peters-Golden M, Kao JY, Markovitz DM. Murine colitis is mediated by vimentin. Sci Rep. 2013;3:1045. Epub 2013/01/11. doi: 10.1038/srep01045. PubMed PMID: 23304436; PMCID: PMC3540396.

39. Erler JT, Weaver VM. Three-dimensional context regulation of metastasis. Clin Exp Metastasis. 2009;26(1):35–49. Epub 2008/09/25. doi: 10.1007/s10585-008-9209-8. PubMed PMID: 18814043; PMCID: PMC2648515.

40. Chandel NS, McClintock DS, Feliciano CE, Wood TM, Melendez JA, Rodriguez AM, Schumacker PT. Reactive oxygen species generated at mitochondrial complex III stabilize hypoxia-inducible factor-1alpha during hypoxia: a mechanism of O2 sensing. J Biol Chem. 2000;275(33):25130–8. Epub 2000/06/02. doi: 10.1074/jbc.M001914200. PubMed PMID: 10833514.

41. Rankin EB, Giaccia AJ. Hypoxic control of metastasis. Science. 2016;352(6282):175–80. Epub 2016/04/29. doi: 10.1126/science.aaf4405. PubMed PMID: 27124451; PMCID: PMC4898055.

42. Martinez-Reyes I, Chandel NS. Mitochondrial TCA cycle metabolites control physiology and disease. Nat Commun. 2020;11(1):102. Epub 2020/01/05. doi: 10.1038/s41467-019-13668-3. PubMed PMID: 31900386; PMCID: PMC6941980.

43. Alvarez-Tejado M, Naranjo-Suarez S, Jimenez C, Carrera AC, Landazuri MO, del Peso L. Hypoxia induces the activation of the phosphatidylinositol 3-kinase/Akt cell survival pathway in PC12 cells: protective role in apoptosis. J Biol Chem. 2001;276(25):22368–74. Epub 2001/04/11. doi: 10.1074/jbc.M011688200. PubMed PMID: 11294857.

44. Romero R, Sayin VI, Davidson SM, Bauer MR, Singh SX, LeBoeuf SE, Karakousi TR, Ellis DC, Bhutkar A, Sanchez-Rivera FJ, Subbaraj L, Martinez B, Bronson RT, Prigge JR, Schmidt EE, Thomas CJ, Goparaju C, Davies A, Dolgalev I, Heguy A, Allaj V, Poirier JT, Moreira AL, Rudin CM, Pass HI, Vander Heiden MG, Jacks T, Papagiannakopoulos T. Keap1 loss promotes Kras-driven lung cancer and results in dependence on glutaminolysis. Nat Med. 2017;23(11):1362–8. Epub 2017/10/03. doi: 10.1038/nm.4407. PubMed PMID: 28967920; PMCID: PMC5677540.

45. Gibbons DL, Lin W, Creighton CJ, Rizvi ZH, Gregory PA, Goodall GJ, Thilaganathan N, Du L, Zhang Y, Pertsemlidis A, Kurie JM. Contextual extracellular cues promote tumor cell EMT and metastasis by regulating miR-200 family expression. Genes Dev. 2009;23(18):2140–51. Epub 2009/09/18. doi: 10.1101/gad.1820209. PubMed PMID: 19759262; PMCID: PMC2751985.

46. Thomas PA, Kirschmann DA, Cerhan JR, Folberg R, Seftor EA, Sellers TA, Hendrix MJ. Association between keratin and vimentin expression, malignant phenotype, and survival in postmenopausal breast cancer patients. Clin Cancer Res. 1999;5(10):2698–703. Epub 1999/10/28. PubMed PMID: 10537332.

47. Peuhu E, Virtakoivu R, Mai A, Warri A, Ivaska J. Epithelial vimentin plays a functional role in mammary gland development. Development. 2017;144(22):4103–13. doi: 10.1242/dev.154229. PubMed PMID: 28947532.

48. Wang Z, Divanyan A, Jourd’heuil FL, Goldman RD, Ridge KM, Jourd’heuil D, Lopez-Soler RI. Vimentin expression is required for the development of EMT-related renal fibrosis following unilateral ureteral obstruction in mice. Am J Physiol Renal Physiol. 2018;315(4):F769–F80. Epub 2018/04/11. doi: 10.1152/ajprenal.00340.2017. PubMed PMID: 29631355; PMCID: PMC6335003.

49. Cheng F, Shen Y, Mohanasundaram P, Lindstrom M, Ivaska J, Ny T, Eriksson JE. Vimentin coordinates fibroblast proliferation and keratinocyte differentiation in wound healing via TGF-beta-Slug signaling. Proceedings of the National Academy of Sciences of the United States of America. 2016;113(30):E4320–7. Epub 2016/07/29. doi: 10.1073/pnas.1519197113. PubMed PMID: 27466403; PMCID: PMC4968728.

50. Ivaska J, Vuoriluoto K, Huovinen T, Izawa I, Inagaki M, Parker PJ. PKCepsilon-mediated phosphorylation of vimentin controls integrin recycling and motility. EMBO J. 2005;24(22):3834–45. Epub 2005/11/05. doi: 10.1038/sj.emboj.7600847. PubMed PMID: 16270034; PMCID: PMC1283946.

51. Thaiparambil JT, Bender L, Ganesh T, Kline E, Patel P, Liu Y, Tighiouart M, Vertino PM, Harvey RD, Garcia A, Marcus AI. Withaferin A inhibits breast cancer invasion and metastasis at sub-cytotoxic doses by inducing vimentin disassembly and serine 56 phosphorylation. Int J Cancer. 2011;129(11):2744–55. Epub 2011/05/04. doi: 10.1002/ijc.25938. PubMed PMID: 21538350.

52. Bargagna-Mohan P, Lei L, Thompson A, Shaw C, Kasahara K, Inagaki M, Mohan R. Vimentin Phosphorylation Underlies Myofibroblast Sensitivity to Withaferin A In Vitro and during Corneal Fibrosis. PLoS One. 2015;10(7):e0133399. Epub 2015/07/18. doi: 10.1371/journal.pone.0133399. PubMed PMID: 26186445; PMCID: PMC4506086.

53. Haversen L, Sundelin JP, Mardinoglu A, Rutberg M, Stahlman M, Wilhelmsson U, Hulten LM, Pekny M, Fogelstrand P, Bentzon JF, Levin M, Boren J. Vimentin deficiency in macrophages induces increased oxidative stress and vascular inflammation but attenuates atherosclerosis in mice. Sci Rep. 2018;8(1):16973. Epub 2018/11/20. doi: 10.1038/s41598-018-34659-2. PubMed PMID: 30451917; PMCID: PMC6242955.

54. Chang HW, Li RN, Wang HR, Liu JR, Tang JY, Huang HW, Chan YH, Yen CY. Withaferin A Induces Oxidative Stress-Mediated Apoptosis and DNA Damage in Oral Cancer Cells. Front Physiol. 2017;8:634. Epub 2017/09/25. doi: 10.3389/fphys.2017.00634. PubMed PMID: 28936177; PMCID: PMC5594071.

55. LeBert D, Squirrell JM, Freisinger C, Rindy J, Golenberg N, Frecentese G, Gibson A, Eliceiri KW, Huttenlocher A. Damage-induced reactive oxygen species regulate vimentin and dynamic collagen-based projections to mediate wound repair. Elife. 2018;7. Epub 2018/01/18. doi: 10.7554/eLife.30703. PubMed PMID: 29336778; PMCID: PMC5790375.

56. Li QF, Spinelli AM, Tang DD. Cdc42GAP, reactive oxygen species, and the vimentin network. Am J Physiol Cell Physiol. 2009;297(2):C299–309. Epub 2009/06/06. doi: 10.1152/ajpcell.00037.2009. PubMed PMID: 19494238; PMCID: PMC2724092.

57. Weinberg F, Hamanaka R, Wheaton WW, Weinberg S, Joseph J, Lopez M, Kalyanaraman B, Mutlu GM, Budinger GR, Chandel NS. Mitochondrial metabolism and ROS generation are essential for Kras-mediated tumorigenicity. Proceedings of the National Academy of Sciences of the United States of America. 2010;107(19):8788–93. doi: 10.1073/pnas.1003428107. PubMed PMID: 20421486; PMCID: PMC2889315.

58. Matveeva EA, Chernoivanenko, I.S. & Minin, A.A. Vimentin intermediate filaments protect mitochondria from oxidative stress. Biochem Moscow Suppl Ser A 2010(4):321–31. doi: https://doi.org/10.1134/S199074781004001X.

59. Matveeva EA, Venkova LS, Chernoivanenko IS, Minin AA. Vimentin is involved in regulation of mitochondrial motility and membrane potential by Rac1. Biol Open. 2015;4(10):1290–7. Epub 2015/09/16. doi: 10.1242/bio.011874. PubMed PMID: 26369929; PMCID: PMC4610213.

60. Nekrasova OE, Mendez MG, Chernoivanenko IS, Tyurin-Kuzmin PA, Kuczmarski ER, Gelfand VI, Goldman RD, Minin AA. Vimentin intermediate filaments modulate the motility of mitochondria. Mol Biol Cell. 2011;22(13):2282–9. Epub 2011/05/13. doi: 10.1091/mbc.E10-09-0766. PubMed PMID: 21562225; PMCID: PMC3128530.

61. Gaude E, Frezza C. Tissue-specific and convergent metabolic transformation of cancer correlates with metastatic potential and patient survival. Nat Commun. 2016;7:13041. Epub 2016/10/11. doi: 10.1038/ncomms13041. PubMed PMID: 27721378; PMCID: PMC5062467.

62. dos Santos G, Rogel MR, Baker MA, Troken JR, Urich D, Morales-Nebreda L, Sennello JA, Kutuzov MA, Sitikov A, Davis JM, Lam AP, Cheresh P, Kamp D, Shumaker DK, Budinger GR, Ridge KM. Vimentin regulates activation of the NLRP3 inflammasome. Nat Commun. 2015;6:6574. doi: 10.1038/ncomms7574. PubMed PMID: 25762200; PMCID: PMC4358756.

63. Huang SH, Chi F, Peng L, Bo T, Zhang B, Liu LQ, Wu X, Mor-Vaknin N, Markovitz DM, Cao H, Zhou YH. Vimentin, a Novel NF-kappaB Regulator, Is Required for Meningitic Escherichia coli K1-Induced Pathogen Invasion and PMN Transmigration across the Blood-Brain Barrier. PLoS One. 2016;11(9):e0162641. Epub 2016/09/23. doi: 10.1371/journal.pone.0162641. PubMed PMID: 27657497; PMCID: PMC5033352.

64. Jiang SX, Slinn J, Aylsworth A, Hou ST. Vimentin participates in microglia activation and neurotoxicity in cerebral ischemia. J Neurochem. 2012;122(4):764–74. Epub 2012/06/12. doi: 10.1111/j.1471-4159.2012.07823.x. PubMed PMID: 22681613.

65. McDonald-Hyman C, Muller JT, Loschi M, Thangavelu G, Saha A, Kumari S, Reichenbach DK, Smith MJ, Zhang G, Koehn BH, Lin J, Mitchell JS, Fife BT, Panoskaltsis-Mortari A, Feser CJ, Kirchmeier AK, Osborn MJ, Hippen KL, Kelekar A, Serody JS, Turka LA, Munn DH, Chi H, Neubert TA, Dustin ML, Blazar BR. The vimentin intermediate filament network restrains regulatory T cell suppression of graft-versus-host disease. J Clin Invest. 2018;128(10):4604–21. doi: 10.1172/JCI95713. PubMed PMID: 30106752; PMCID: PMC6159973.

66. Richardson AM, Havel LS, Koyen AE, Konen JM, Shupe J, Wiles WGt, Martin WD, Grossniklaus HE, Sica G, Gilbert-Ross M, Marcus AI. Vimentin Is Required for Lung Adenocarcinoma Metastasis via Heterotypic Tumor Cell-Cancer-Associated Fibroblast Interactions during Collective Invasion. Clin Cancer Res. 2018;24(2):420–32. doi: 10.1158/1078-0432.CCR-17-1776. PubMed PMID: 29208669; PMCID: PMC5771825.

67. Karki R, Kanneganti TD. Diverging inflammasome signals in tumorigenesis and potential targeting. Nat Rev Cancer. 2019;19(4):197–214. Epub 2019/03/08. doi: 10.1038/s41568-019-0123-y. PubMed PMID: 30842595; PMCID: PMC6953422.

68. Guo B, Fu S, Zhang J, Liu B, Li Z. Targeting inflammasome/IL-1 pathways for cancer immunotherapy. Sci Rep. 2016;6:36107. Epub 2016/10/28. doi: 10.1038/srep36107. PubMed PMID: 27786298; PMCID: PMC5082376.

69. Al-Saad S, Al-Shibli K, Donnem T, Persson M, Bremnes RM, Busund LT. The prognostic impact of NF-kappaB p105, vimentin, E-cadherin and Par6 expression in epithelial and stromal compartment in non-small-cell lung cancer. Br J Cancer. 2008;99(9):1476–83. Epub 2008/10/16. doi: 10.1038/sj.bjc.6604713. PubMed PMID: 18854838; PMCID: PMC2579693.

70. Lanier MH, Kim T, Cooper JA. CARMIL2 is a novel molecular connection between vimentin and actin essential for cell migration and invadopodia formation. Mol Biol Cell. 2015;26(25):4577–88. Epub 2015/10/16. doi: 10.1091/mbc.E15-08-0552. PubMed PMID: 26466680; PMCID: PMC4678016.

71. Challa AA, Stefanovic B. A novel role of vimentin filaments: binding and stabilization of collagen mRNAs. Mol Cell Biol. 2011;31(18):3773–89. doi: 10.1128/MCB.05263-11. PubMed PMID: 21746880; PMCID: PMC3165730.

72. Gan Z, Ding L, Burckhardt CJ, Lowery J, Zaritsky A, Sitterley K, Mota A, Costigliola N, Starker CG, Voytas DF, Tytell J, Goldman RD, Danuser G. Vimentin Intermediate Filaments Template Microtubule Networks to Enhance Persistence in Cell Polarity and Directed Migration. Cell Syst. 2016;3(3):252–63 e8. Epub 2016/09/27. doi: 10.1016/j.cels.2016.08.007. PubMed PMID: 27667364; PMCID: PMC5055390.

73. Costigliola N, Ding L, Burckhardt CJ, Han SJ, Gutierrez E, Mota A, Groisman A, Mitchison TJ, Danuser G. Vimentin fibers orient traction stress. Proceedings of the National Academy of Sciences of the United States of America. 2017;114(20):5195–200. Epub 2017/05/04. doi: 10.1073/pnas.1614610114. PubMed PMID: 28465431; PMCID: PMC5441818.

74. Eden E, Lipson D, Yogev S, Yakhini Z. Discovering motifs in ranked lists of DNA sequences. PLoS Comput Biol. 2007;3(3):e39. Epub 2007/03/27. doi: 10.1371/journal.pcbi.0030039. PubMed PMID: 17381235; PMCID: PMC1829477.

75. Eden E, Navon R, Steinfeld I, Lipson D, Yakhini Z. GOrilla: a tool for discovery and visualization of enriched GO terms in ranked gene lists. BMC Bioinformatics. 2009;10:48. doi: 10.1186/1471-2105-10-48. PubMed PMID: 19192299; PMCID: PMC2644678.

76. Zhang L, Lee NJ, Nguyen AD, Enriquez RF, Riepler SJ, Stehrer B, Yulyaningsih E, Lin S, Shi YC, Baldock PA, Herzog H, Sainsbury A. Additive actions of the cannabinoid and neuropeptide Y systems on adiposity and lipid oxidation. Diabetes Obes Metab. 2010;12(7):591–603. Epub 2010/07/02. doi: 10.1111/j.1463-1326.2009.01193.x. PubMed PMID: 20590734.

77. Gilbert-Ross M, Konen J, Koo J, Shupe J, Robinson BS, Wiles WGt, Huang C, Martin WD, Behera M, Smith GH, Hill CE, Rossi MR, Sica GL, Rupji M, Chen Z, Kowalski J, Kasinski AL, Ramalingam SS, Fu H, Khuri FR, Zhou W, Marcus AI. Targeting adhesion signaling in KRAS, LKB1 mutant lung adenocarcinoma. JCI Insight. 2017;2(5):e90487. Epub 2017/03/16. doi: 10.1172/jci.insight.90487. PubMed PMID: 28289710; PMCID: PMC5333956 exists.

78. Chong J, Soufan O, Li C, Caraus I, Li S, Bourque G, Wishart DS, Xia J. MetaboAnalyst 4.0: towards more transparent and integrative metabolomics analysis. Nucleic Acids Res. 2018;46(W1):W486–W94. Epub 2018/05/16. doi: 10.1093/nar/gky310. PubMed PMID: 29762782; PMCID: PMC6030889.

