## Supplemental Figures for "Vimentin is Required for Tumor Progression and Metastasis in a Mouse Model of Non-Small Cell Lung Cancer"

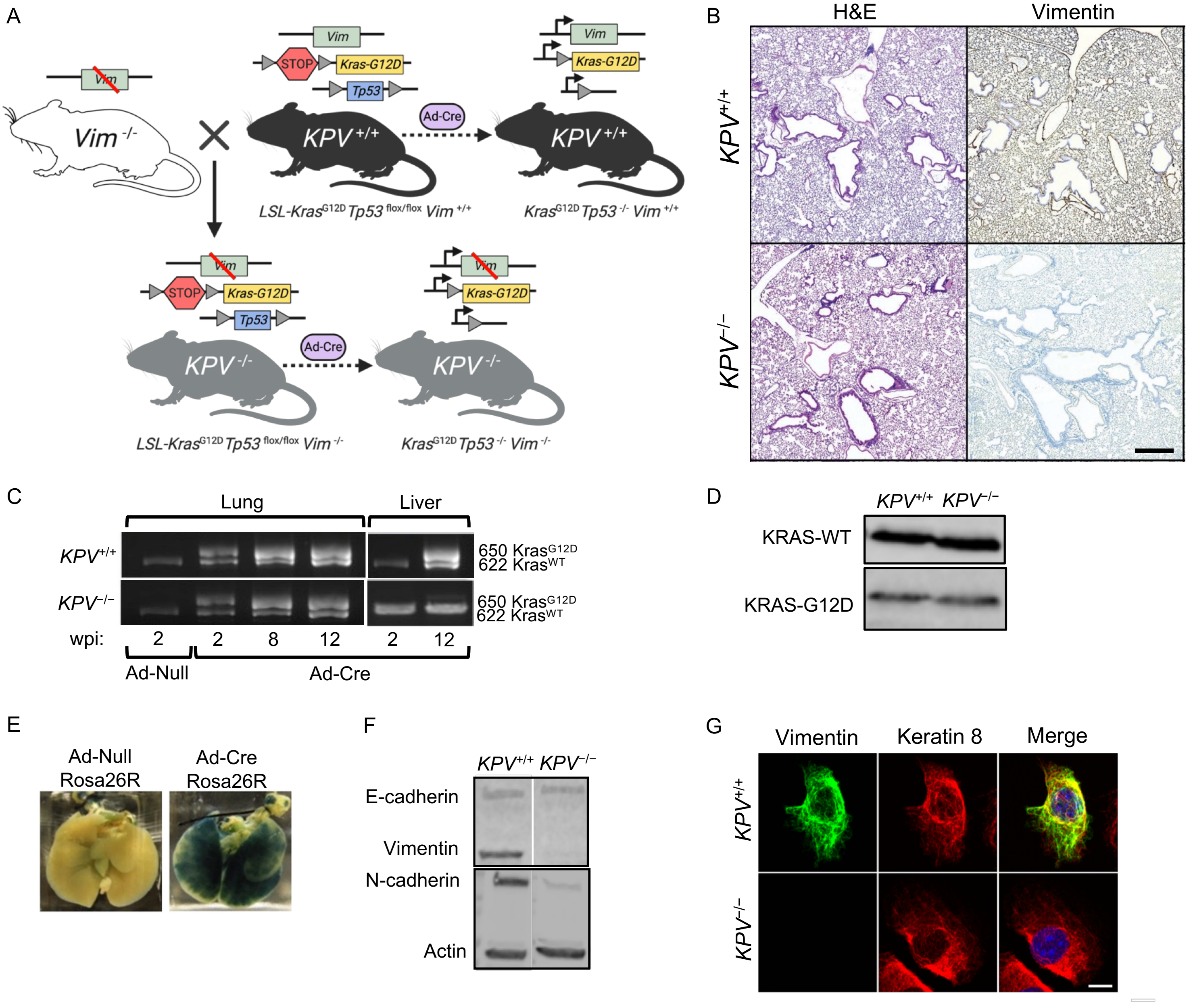


**Supplemental Figure 1.** (**A**) Experimental design. *LSL-Kras*$^{G12D/+}$*Tp53*$^{flox/flox}$ (*KPV*$^{+/+}$) mice were crossed with vimentin-knockout (*Vim*$^{-/-}$) mice. *KPV*$^{+/+}$ and *KPV*$^{-/-}$ mice were administered adenoviral Cre recombinase (Ad-Cre) which resulted in gene recombination at LoxP sites. As a control, null adenovirus (Ad-null) was administered to an independent cohort of mice. (**B**) Lungs were isolated from Ad-null-treated *KPV*$^{+/+}$ and *KPV*$^{-/-}$ mice, fixed, sectioned, and subjected to H&E staining and vimentin immunohistochemical staining. Positive vimentin staining is brown, and nuclei are blue. Scale bar: 200 µm. (**C**) *KPV*$^{+/+}$ and *KPV*$^{-/-}$ mice were administered Ad-Null or Ad-Cre. Lungs and livers were harvested at 2, 8, and 12 weeks following adenoviral infection. DNA was isolated from the tissue and PCR was performed to evaluate the presence of the wild-type (WT) and mutant (G12D) *Kras* transcript. (**D**) Tumor cells were isolated from Ad-Cre-infected mice at 6 weeks post-infection. A Western blot was performed on *KPV*$^{+/+}$ and *KPV*$^{-/-}$ whole cell lysates to detect WT and G12D-mutant KRAS. (**E**) *Rosa26-LSL-LacZ* reporter mice were administered Ad-Null or Ad-Cre, and β-galactosidase staining was performed on whole lung samples; positive staining appears blue. (**F**) *KPV*$^{+/+}$ and *KPV*$^{-/-}$ cell lysates were subjected to a Western blot for detection of E-cadherin, vimentin, N-cadherin, and actin. (**G**) *KPV*$^{+/+}$ and *KPV*$^{-/-}$ cells were stained for vimentin (*green*), for keratin 8 (*red*), and with Hoechst nuclear dye (*blue*). Scale bar: 10 µm.

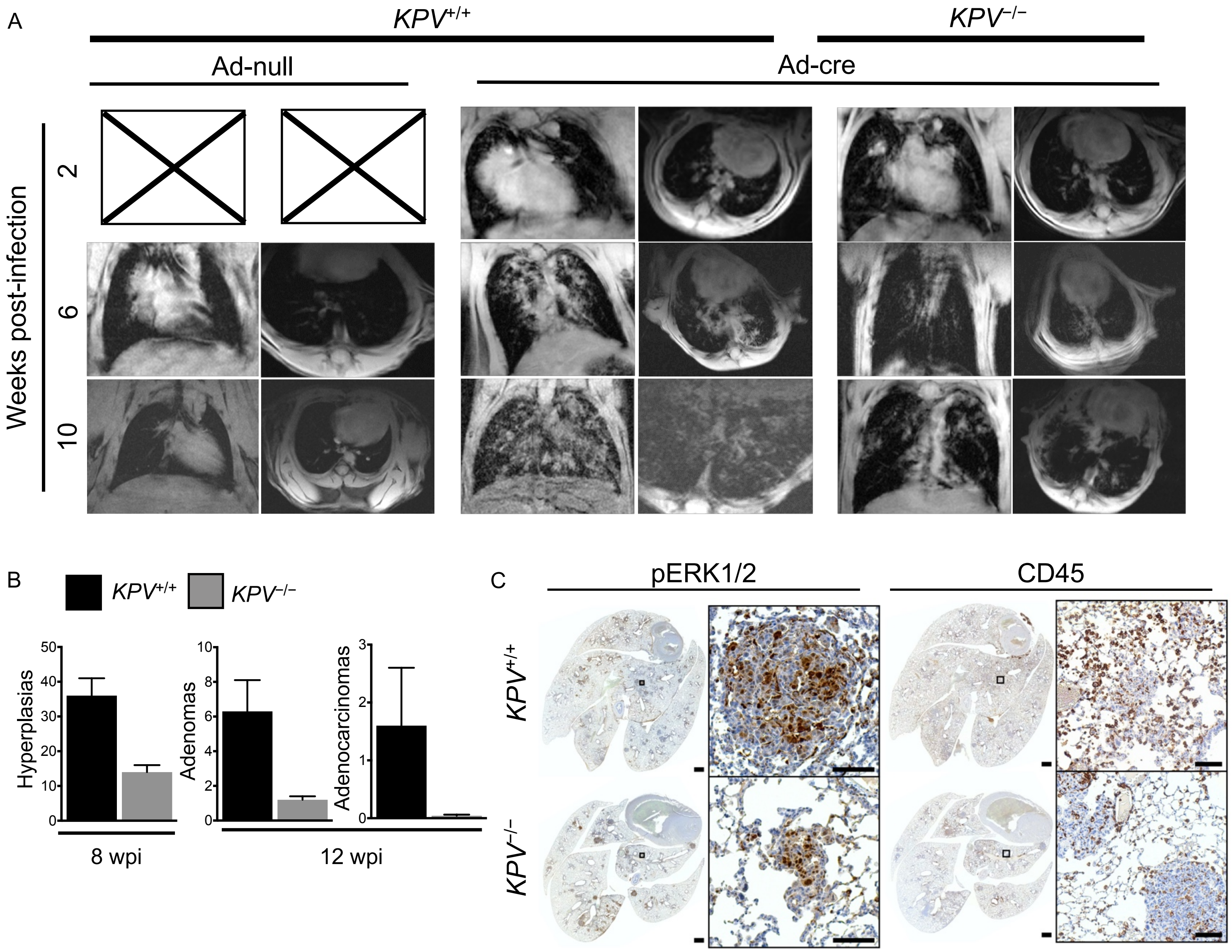


**Supplemental Figure 2.** (**A**) *KPV*$^{+/+}$ and *KPV*$^{-/-}$ mice were infected with null or Cre recombinase adenovirus (Ad-Null and Ad-Cre, respectively) and were imaged at 2, 6, and 10 weeks post-infection (wpi). Representative MRI coronal (*left*) and transverse (*right*) images are shown. (**B**) H&E-stained lung sections from 8 or 12 wpi were evaluated for tumor grade by a pathologist. (**C**) Lungs were harvested from *KPV*$^{+/+}$ and *KPV*$^{-/-}$ mice at 7 wpi, fixed, sectioned, and subjected to immunohistochemistry with antibodies against CD45 and phospho-ERK1/2. Scale bar: 1 mm (whole lung), 100 µm (insets).

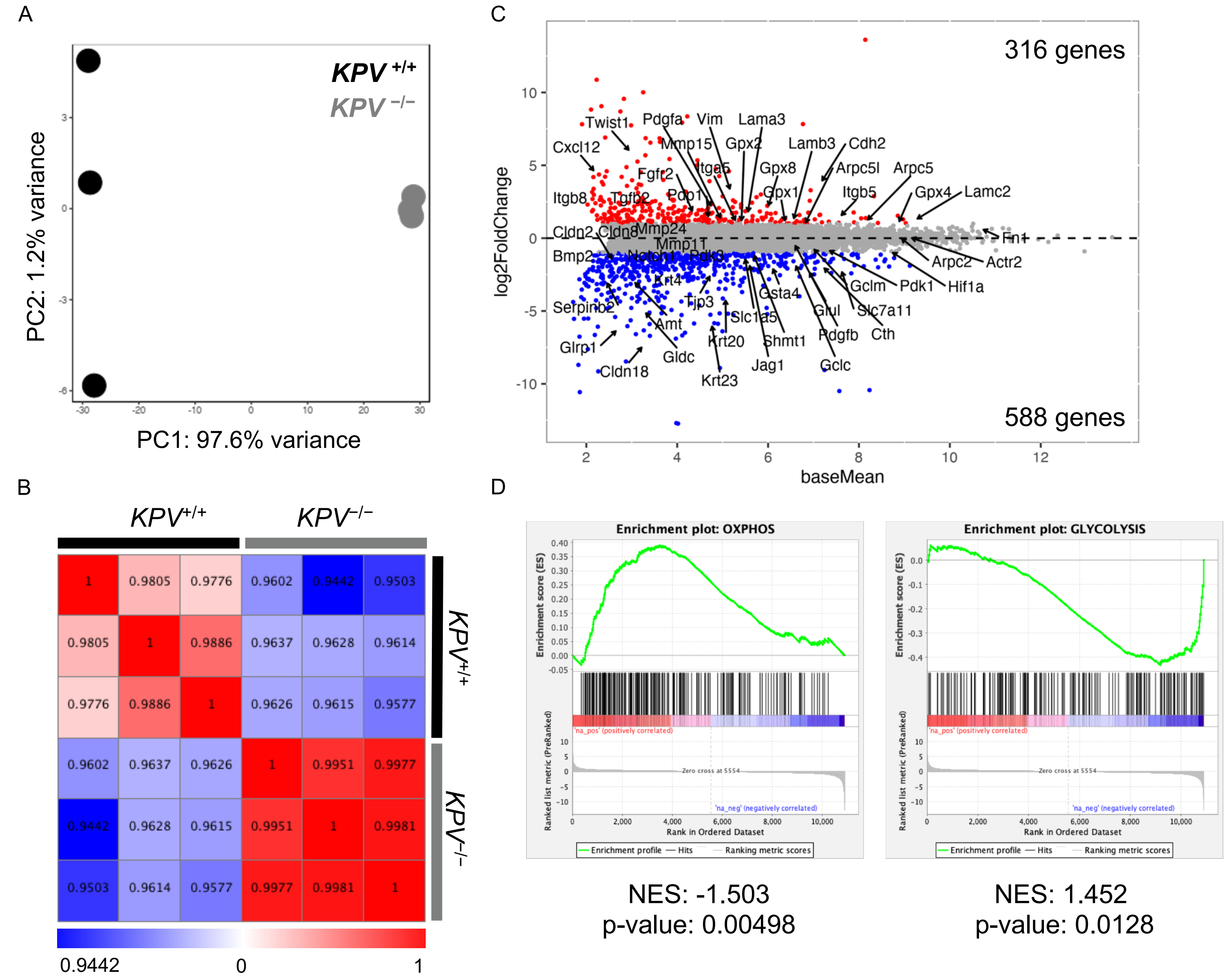


**Supplemental Figure 3.** Messenger RNA from *KPV*$^{+/+}$ and *KPV*$^{-/-}$ cell lysates was quantified via RNA-sequencing. (**A**) Principal component analysis (PCA) plot with each point representing one replicate (*black, KPV*$^{+/+}$; *grey, KPV*$^{-/-}$). (**B**) Pearson's correlation plot. The correlation coefficient for each comparison is shown. (**C**) MA plot. Genes of interest are indicated. (**D**) Gene set enrichment analysis (GSEA) plots. MSigDB hallmark pathways "OXPHOS" (oxidative phosphorylation) and "GLYCOLYSIS" were significantly enriched in *KPV*$^{+/+}$ and *KPV*$^{-/-}$ cells, respectively ($p<0.05$). Normalized enrichment score (NES) and p-value are shown below each plot.

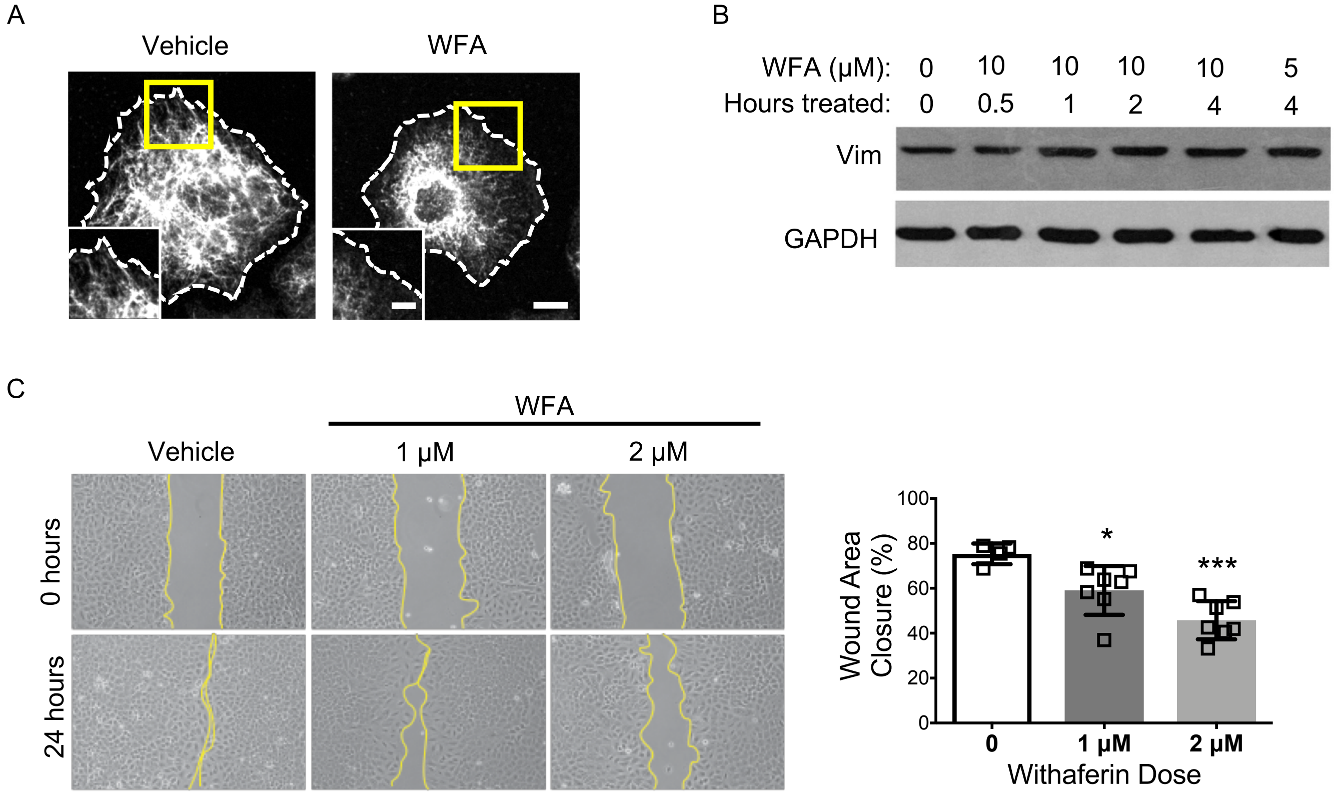


**Supplemental Figure 4.** (**A**) A549 cells were treated with 2 µM withaferin A (WFA) for 1 hour. Cells were fixed and stained for vimentin (white). A phase contrast image was used to identify cell borders (dashed line). Scale bar: 10 µm, 5 µm *(inset*). (**B**) A549 cells were treated with WFA for the indicated dose and time. A Western blot is shown; vimentin and GAPDH antibodies were used to probe for these proteins. (**C**) A549 cells were treated with DMSO control or with 1 or 2 µM WFA and were subjected to a scratch wound assay. After 24 hours, wound closure was assessed. Representative images (*left*) and quantitation (*right*) are shown. Data were compared to the vehicle control using an unpaired, two-tailed t-test (*$p<0.05$; ***$p<0.001$).

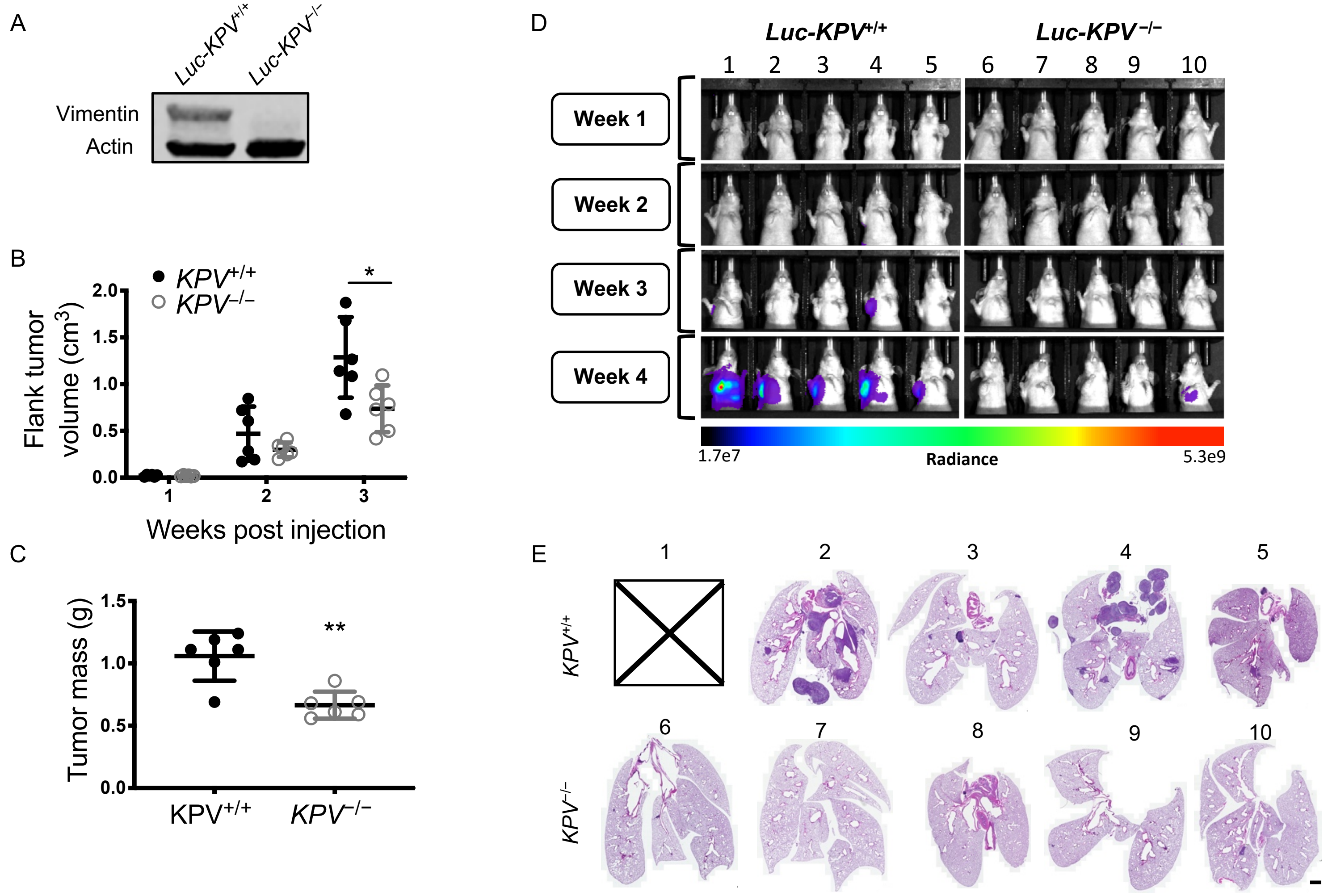


**Supplemental Figure 5.** (**A**) Luciferase-tagged *KPV*$^{+/+}$ (*Luc-KPV*$^{+/+}$) cells were treated with CRISPR-Cas9 to knock out vimentin (*Luc-KPV*$^{-/-}$). Cells were subjected to a Western blot and probed for vimentin and actin. (**B**) Nude mice were injected subcutaneously with *Luc-KPV*$^{+/+}$ or *Luc-KPV*$^{-/-}$ cells. Flank tumor volume was measured with calipers each week. Volume was calculated using the formula Volume=(length$^2$ x width)/2. (**C**) At week 3, tumors were removed and weighed. For **B-C**, data were subjected to an unpaired, two-tailed t-test. (*$p<0.05$; **$p<0.01$). (**D**) IVIS images of mouse thoracic region. At weeks 3 and 4, primary tumors were covered to limit signal bleed-through. (**E**) Mice were sacrificed at week 4. Lungs were harvested, fixed, sectioned, and stained with H&E. Each numbered lung corresponds with the numbered mouse in **D**. Mouse 1 died before lungs could be harvested. Scale bar: 1 mm.

| Antibody Target | Company | Host Species | Clone | Catalog Number | Use |
|---|---|---|---|---|---|
| Actin | Santa Cruz Biotechnology | Mouse | C-2 | sc-8432 | WB |
| Actin | Santa Cruz Biotechnology | Goat | C-11 | sc-1615 | WB |
| Akt1 | Cell Signaling Technology | Rabbit | C73H10 | 2938 | WB |
| Akt2 | Cell Signaling Technology | Rabbit | D6G4 | 3063 | WB |
| CD45 | Abcam | Rabbit | polyclonal | ab10558 | IHC |
| E-cadherin | Santa Cruz Biotechnology | Rabbit | H-108 | sc-7870 | WB |
| GAPDH | Cell Signaling Technology | Rabbit | 14C10 | 2118 | WB |
| Keratin | Fitzgerald | Mouse | Ks8.7 | 10R-C177ax | IF |
| Ki67 | Abcam | Rabbit | polyclonal | ab66155 | IHC |
| Kras | Abcam | Rabbit | polyclonal | ab180772 | WB |
| N-cadherin | Calbiochem | Rabbit | polyclonal | 205606 | WB |
| pAkt | Cell Signaling Technology | Mouse | 587F11 | 4051 | WB |
| pan-Akt | Cell Signaling Technology | Rabbit | 11E7 | 4685 | WB |
| pERK1/2 | Cell Signaling Technology | Rabbit | D13.14.4E | 4370 | IHC |
| Ras(G12D) | NewEast Biosciences | Mouse | | 26036 | WB |
| TTF-1 | Abcam | Rabbit | EPR8190 | ab133638 | IHC |
| Vimentin | Abcam | Rabbit | EPR3776 | ab92547 | IHC, IF (Fig. S1G) |
| Vimentin | Biolegend | Chicken | Poly29191 | 919101 | IF (Fig. 3E) |
| Vimentin | Cell Signaling Technology | Rabbit | R28 | 3932 | IF (Fig. 6B) |
| Vimentin | Sigma-Aldrich | Mouse | V9 | V6630 | WB (Fig. 6C, F-H) |

**Supplemental Table 1.** List of antibodies used. For proteins with multiple antibodies used, the figures in which they are used are indicated. If no figure is indicated, the antibody was used for all instances in which that protein was detected. **WB**=Western blot; **IHC**=immunohistochemistry; **IF**=immunofluorescence.
